# Optimizing plant species selection for automated monitoring of plant-pollinator interactions

**DOI:** 10.64898/2026.09.22.753407

**Authors:** Yuping Zhong, Jose B. Lanuza, Jonna M. Heuschele, Will Glenny, Demetra Rakosy, Tiffany M. Knight

## Abstract

Automated monitoring camera systems offer an efficient approach for quantifying pollinator biodiversity and plant-pollinator interactions, but financial and logistical constraints limit the number of flowering plant species that can be monitored. Using a large European database of plant-pollinator networks, we evaluated whether plant subsampling can capture key metrics of interest (i.e. pollinator richness and network structure). We compared abundance-based, flower-shape-informed, phylogenetically informed and random plant sampling. Abundance-based sampling consistently outperformed random sampling, while adding flower shape provided little additional benefit and phylogenetic selection performed similarly to random sampling. Overall, monitoring 12-15 flowering species was sufficient to characterize key network properties across varying community sizes. Performance of abundance-based plant sampling declined in species-rich communities and when rare but highly attractive plants were present in the community, while a specific subset of pollinator species was consistently missed even by the best sampling strategy. Based on their relative contribution to interactions within each network, only 12.6% of pollinator species accounted for 95% of interactions, suggesting that AI classifiers could prioritize a relatively small subset of species. Our results provide practical guidance for designing efficient camera-based pollinator monitoring schemes at large spatial scales.

## Introduction

Non-lethal automated monitoring of insect pollinators using cameras and machine learning is emerging as a rapid and cost-effective approach for large-scale pollinator monitoring [1]. Cameras aimed at flowers can capture images of visiting pollinators [2–4] and machine learning can identify these pollinators [5,6]. Because automated monitoring facilitates standardized and efficient data collection, it has the potential to capture detailed information on pollinator diversity and plant-pollinator interactions across space and time. However, logistical and financial constraints make it impractical to deploy cameras on all flowering plant species within a community, particularly in species-rich systems where many plant species may flower simultaneously. As a result, optimizing plant species selection is essential for the success of large-scale automated monitoring. Empirical guidance for selecting the optimal number and composition of plant species for monitoring is currently lacking.

Species abundances and traits are key determinants of species interactions [7,8], providing a basis for optimizing plant species selection. More abundant plant species are visited by a wider range of pollinator species [9–11], while floral traits such as floral morphology mediate both flower attractiveness [12–14] and access to floral rewards [15]. Both abundance and flower shape are relatively easy to characterize, making them practical criteria for plant selection in monitoring programs. However, the usefulness of broad flower-shape categories may depend on how consistently they predict pollinator assemblages across plant species and communities. Sampling effectiveness may further depend on how interactions are distributed among species within communities. For example, in more nested networks, less-connected species tend to interact with subsets of the partners of highly connected species, which may increase the redundancy of information captured during plant sampling. Moreover, plant attractiveness to pollinators is shaped by additional floral traits such as flower color, nectar reward quality, and scent [12,16], which may cause certain plant species to attract disproportionately more pollinators than their abundance or morphology alone would predict. Because the evolutionary history of a species can act as a surrogate for multiple unmeasured floral traits, phylogenetically informed sampling may help overcome this limitation, although its implementation requires a local plant inventory and an available phylogeny [17] . It remains unclear how these plant characteristics and community-level interaction properties influence the ability of subsampling strategies to capture pollinator diversity and network structure.

Europe offers considerable potential for automated pollinator monitoring because plant communities are already surveyed annually at hundreds of sites through the European Long-Term Ecological Research Infrastructure (eLTER) [18]. These surveys provide an opportunity to integrate camera-based monitoring of pollinators and plant-pollinator interactions into existing monitoring programs, facilitating standardized large-scale monitoring. Likewise, while thousands of insect species are potential pollinators in Europe, a subset of common species are involved in most of the interactions [19]. This pattern suggests that prioritizing frequently occurring species for image-based classification could improve both monitoring efficiency and model performance. Existing image databases are often taxonomically biased [20–22], highlighting the need to identify priority pollinator taxa for classifier development.

Here, we used the European Database of Plant-Pollinator Networks (EuPPollNet), a continental-scale dataset of plant-pollinator interactions across Europe, to evaluate the performance of random and ecologically informed sampling approaches. We compared three ecologically-informed sampling approaches against a random sampling benchmark: selecting the most abundant flowering plant species in the community, selecting abundant species across different floral morphologies, and selecting a set of species that maximized phylogenetic distances within the network. Each sampling approach was evaluated at different levels of sampling effort (three, five and ten flowering plant species). We specifically assessed how well these sampling strategies capture pollinator richness, plant-pollinator interactions and network structure by comparing subsampled with the original traditionally sampled networks. We also assessed whether the performance of each sampling strategy depended on the inclusion of highly attractive plant species that support disproportionately high numbers and diversity of interactions. In addition, we identified pollinator taxa consistently missed by the best-performing strategy to highlight detection gaps and identified priority pollinator taxa for AI-based classification in European monitoring programs. Together, these analyses provide an ecological basis for optimizing automated pollinator monitoring, improving our understanding of pollinator communities and interactions, and informing conservation efforts.

## Results

### Pollinator richness across sampling strategies

Both abundance-based and flower-shape abundance-based subsampling outperformed random sub-sampling at the same sampling effort, while phylogenetic distance-based subsampling showed no improvement over the random benchmark (**Fig. 1**, Supplementary **Table S1**). The two approaches performed similarly when selecting three species, while abundance-based subsampling captured slightly more pollinator richness when selecting five species (Holm-adjusted P = 0.036). These patterns were consistent across major pollinator orders (Supplementary **Fig. S1**). On average, sampling 40% of flowering plant species per network was sufficient to capture 80% of pollinator richness with the best strategy, with additional sampling yielding progressively smaller gains in pollinator richness (Supplementary **Fig.S2**).

**Figure 1.**
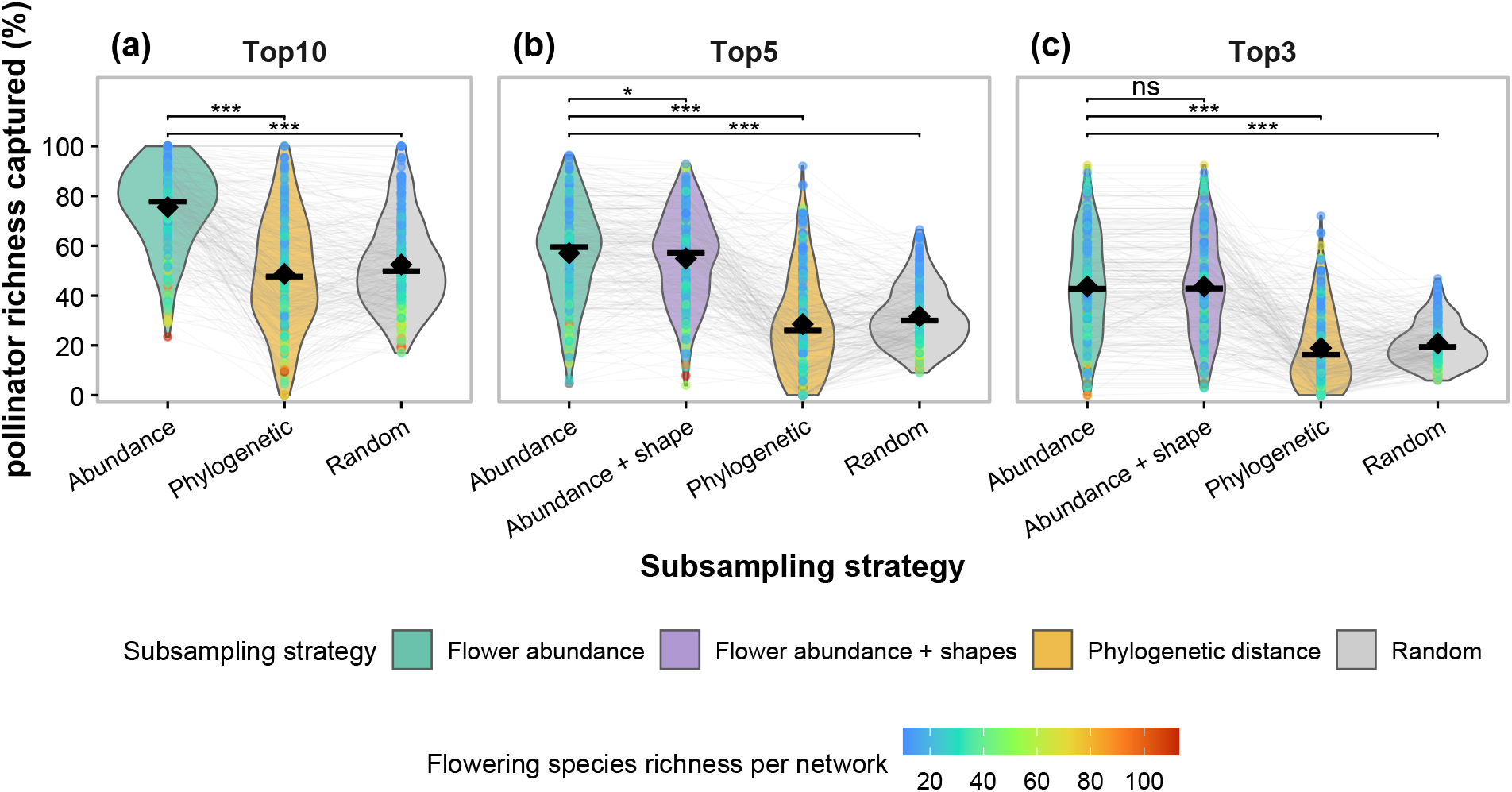
Percentage of pollinator richness captured under different plant sampling strategies. Panels show sampling of ten, five, and three plant species, respectively. Violin colors indicate the sampling strategy: abundance-based (green), flower-shape abundance-based (purple), phylogenetic distance-based (yellow), and random sampling (gray). Points represent individual networks, with point color indicating the total number of flowering plant species in each network. Black diamonds indicate means, and horizontal black bars indicate medians. Flower-shape abundance-based sampling was not considered when sampling ten species because too few networks contained ten or more distinct flower-shape categories. Asterisks indicate unadjusted P values from two-sided paired Wilcoxon tests (*P < 0*.*05*, ***P < 0*.*01***, P < 0.001); ns, not significant.

### Network structure across sampling strategies

The similarity between subsampled and complete networks depended on sampling effort, community richness, and interaction structure. Among the three sampling strategies implemented, sampling more species (i.e., ten species) yielded stronger associations, with connectance showing more sensitivity to varying sampling effort than nestedness and selectivity (Supplementary **Fig. S3**). Extending the comparison between abundance-based and random subsampling across a broader range of sampling efforts revealed that the strength of the association increased with the number of plant species sampled. Strong association (ρ ≥ 0.8) were reached at lower sampling efforts under abundance-based than random subsampling (**Fig. 2**). Specifically, abundance-based subsampling required 15 abundant plant species for connectance, 13 for NODF and 12 for selectivity whereas random selection required 22, 27 and 43 plant species, respectively, to reach the strong association threshold. A complementary proportional-sampling analysis showed that capturing at least 80% of pollinator richness required, on average, approximately 30-45% of flowering plant species under abundance-based selection, compared with approximately 45-72% under random selection (Supplementary **Fig. S4**)

**Figure 2.**
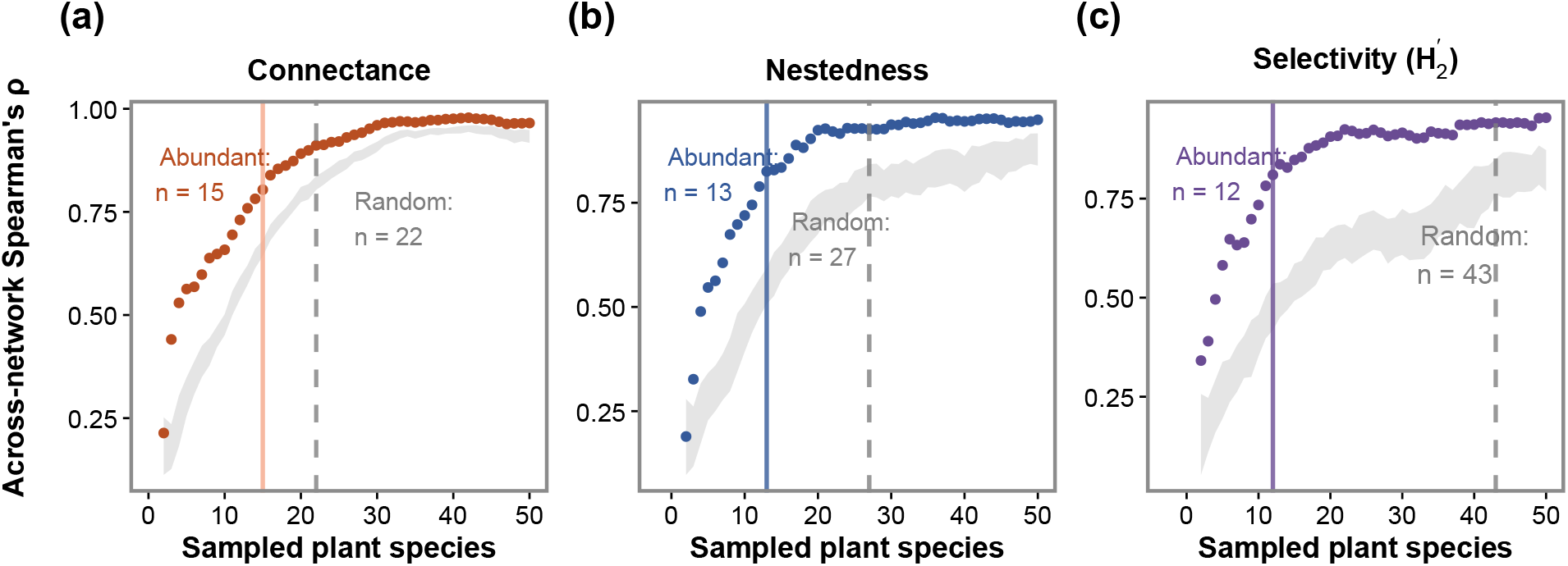
Across-network Spearman’s rank correlation () between subsampled and complete network metrics as a function of the number of sampled plant species. (a) Connectance, (b) nestedness (NODF), and (c) selectivity (H). Points represent subnetworks generated by sequentially adding the most abundant plant species. The gray shaded area represents ±1 SD around the mean Spearman’s across 50 random plant-selection replicates. Vertical lines indicate the minimum observed number of plant species at which first reaches the predefined threshold of 0.8 for the abundance-based and random-selection approaches.

### Factors associated with plant sampling effectiveness

Abundance-based sampling was the most effective strategy in capturing pollinator diversity and network structure. However, its effectiveness varied substantially among networks, largely depending on community richness and interaction structure. As plant and pollinator richness increased, the percentage of pollinator richness captured at a given sampling effort decreased (**Fig. 3a**), while greater nestedness of the original network increased the effectiveness of abundance-based sampling in capturing pollinator richness (**Fig. 3b**).

**Figure 3.**
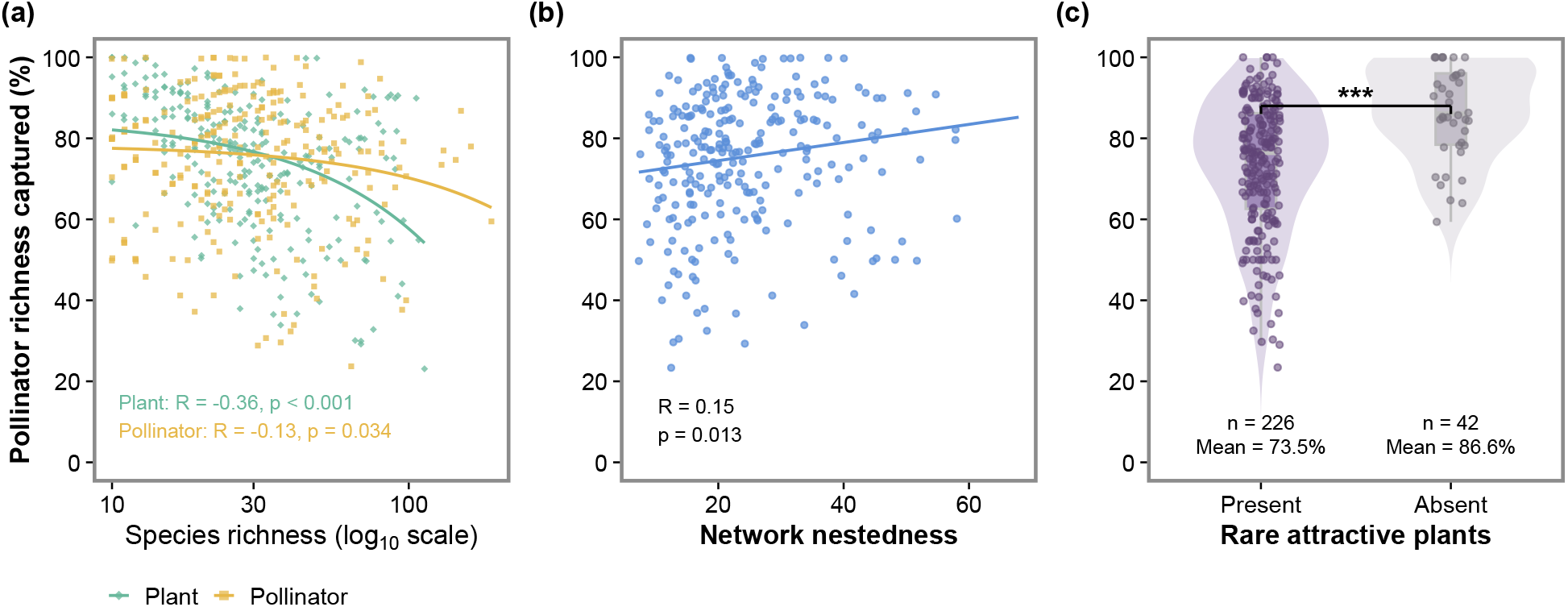
Factors associated with the effectiveness of the abundance-based sampling strategy. (a) Relationships between percentage of pollinator richness captured by sampling the ten most abundant plant species and total plant richness and total pollinator richness per network. The richness axis is displayed on a log_10_ scale, whereas Pearson’s correlation coefficients were calculated using richness on its original scale. (b) Relationship between the percentage of pollinator richness captured by the ten most abundant plant species and nestedness of the complete network. Pearson correlation coefficients (R) and its corresponding p-value (p) are shown. Lines represent linear model fits for visualization purposes. (c) Percentage of pollinator richness captured when sampling the ten most abundant plant species. “Present” indicates that at least one of the 47 highly attractive plant species occurred in the network but was not among the 10 most abundant species, whereas “Absent” indicates that they were either absent from the network or captured by the top 10 most abundant species.

The presence and abundance of highly attractive plant species also had a substantial impact on sampling performance. Pollinator richness was captured more effectively when highly attractive plant species were adequately represented in the sampled plants. This occurred when these species were either absent from the community or sufficiently abundant to be included among the sampled plants. In contrast, pollinator richness capture was lower when at least one highly attractive species was present but too rare to be included among the ten most abundant plants (86.55 ± 11.68% vs. 73.45 ± 16.78%; Wilcoxon rank-sum test, *P* = 1.5 × 10 ; **Fig. 3c**).

The composition of pollinator genera visiting flowers significantly differed across flower shape categories after accounting for study-level variation and restricting permutations within networks (PERMANOVA, F = 20.70, R^2^ = 0.025, P = 0.001; **Fig. 4a**), but flower shape explained only 2.5% of the variation in pollinator community composition. Significant dispersion differences among flower shapes (PERMDISP: F = 125.77, P = 0.001; Supplementary **Fig. S5**) indicated the PERMANOVA effect may partly reflect differences in within-group dispersion rather than differences in community centroids alone. Pollinator assemblages also varied substantially among plant species sharing the same flower shape within the same network (**Fig. 4b**). Mean within-network Sørensen dissimilarity differed among flower-shape categories (linear mixed-effects model: *F* = 10.27, *P* < 0.001), but was generally high across categories. In particular, flag, lip, and bell flowers exhibited significantly lower within-network dissimilarity than open disk flowers, stalk disk flowers and flower heads, indicating more homogeneous pollinator communities associated with plant species belonging to the former categories.

**Figure 4.**
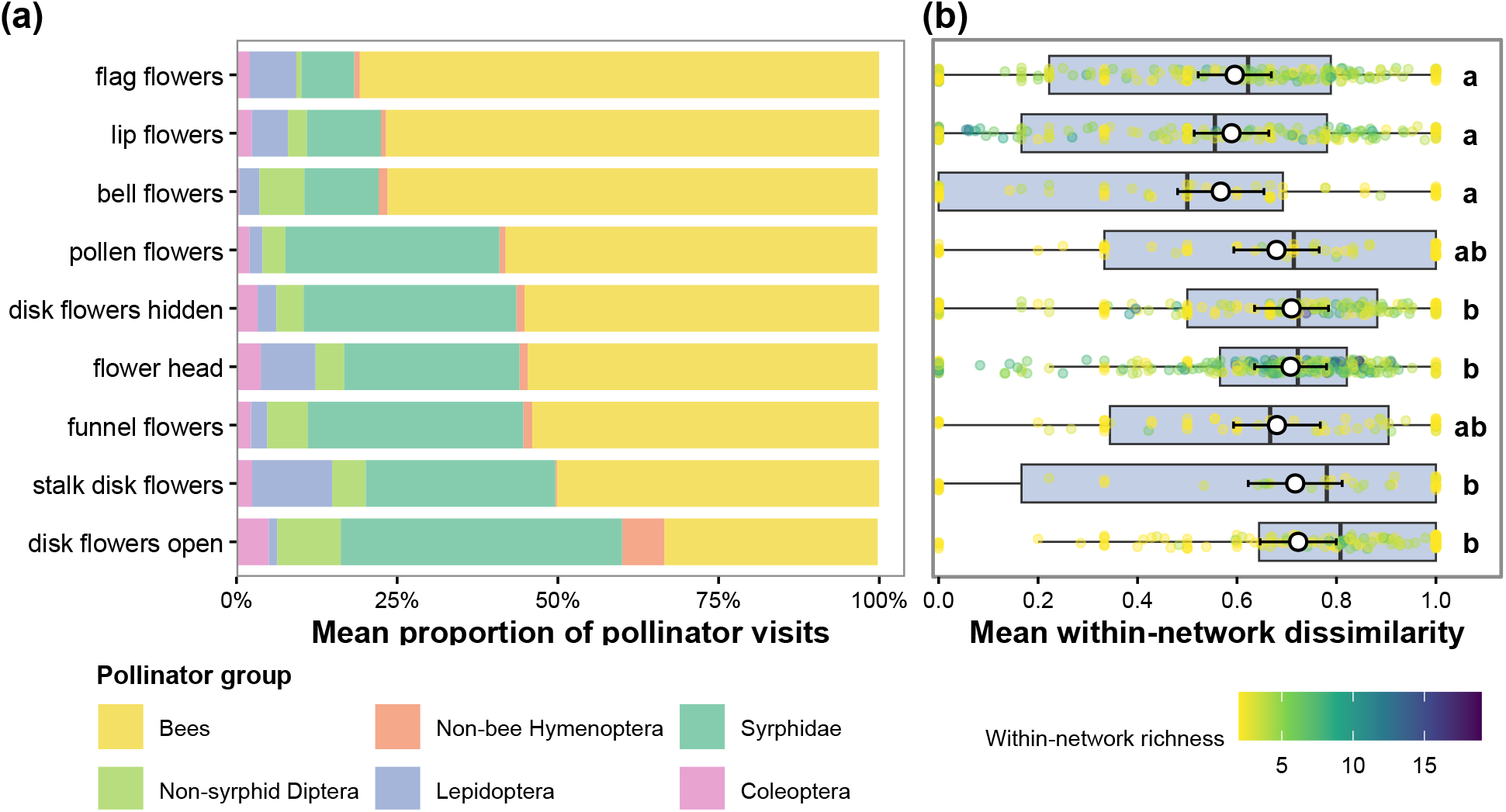
Variation in pollinator community composition and its sources across flower shapes. (a) Pollinator functional-group composition and sampling representation across flower shapes. Stacked bars show the mean relative proportion of visits by pollinator functional groups for each flower-shape category across networks. *Apis* records were excluded from the functional-group panel because of their broad distribution and high abundance, and trap flowers because they received fewer than four visits. (b) Within-network variation in pollinator assemblage composition among plant species sharing the same flower-shape category. Each colored point represents one flower-shape × network combination and shows the mean pairwise Sørensen dissimilarity among its constituent plant species. Point color indicates the number of plant species included in that combination. Boxplots show the observed distributions. White points and error bars show estimated marginal means and 95% confidence intervals from the linear mixed-effects model. Different letters indicate significant pairwise differences following Sidak adjustment.

### Target pollinator species for AI classification

We found that monitoring only the most abundant plant species missed 21.6-26.7% of species-by-network records across the four focal insect orders, with the highest pooled miss rate in Lepidoptera (26.7%), followed by Hymenoptera (26.3%), Diptera (25.1%), and Coleoptera (21.6%). Miss rates also varied among families (Supplementary **Fig. S6**). For example, Pieridae had a miss rate of 35.7% (35/98), while Andrenidae, Megachilidae, and Colletidae each had rates of approximately 31%, compared with 22.9% for Apidae. Apidae species differed markedly in their susceptibility to being missed by abundance-based plant sampling. Missed rates ranged from 5% to 86%, with several *Bombus* species frequently underrepresented (**Table S2**). This pattern was not restricted to infrequently recorded species: some widespread bumblebees were missed from a substantial number of networks, including *Bombus pratorum* (31 of 102 networks) and *B. hortorum* (29 of 112), whereas *Apis mellifera* was missed from only 11 of 215 networks (5%).

Across the entire dataset of 1,630 European plant-pollinator networks, 2,223 insect species were recorded, but only 279 species (12.6% of the total) accounted for 95% of the cumulative mean relative interaction contribution across networks (**Fig. 5**, Supplementary **Fig. S7;** Supplementary **Dataset S1**).

**Figure 5.**
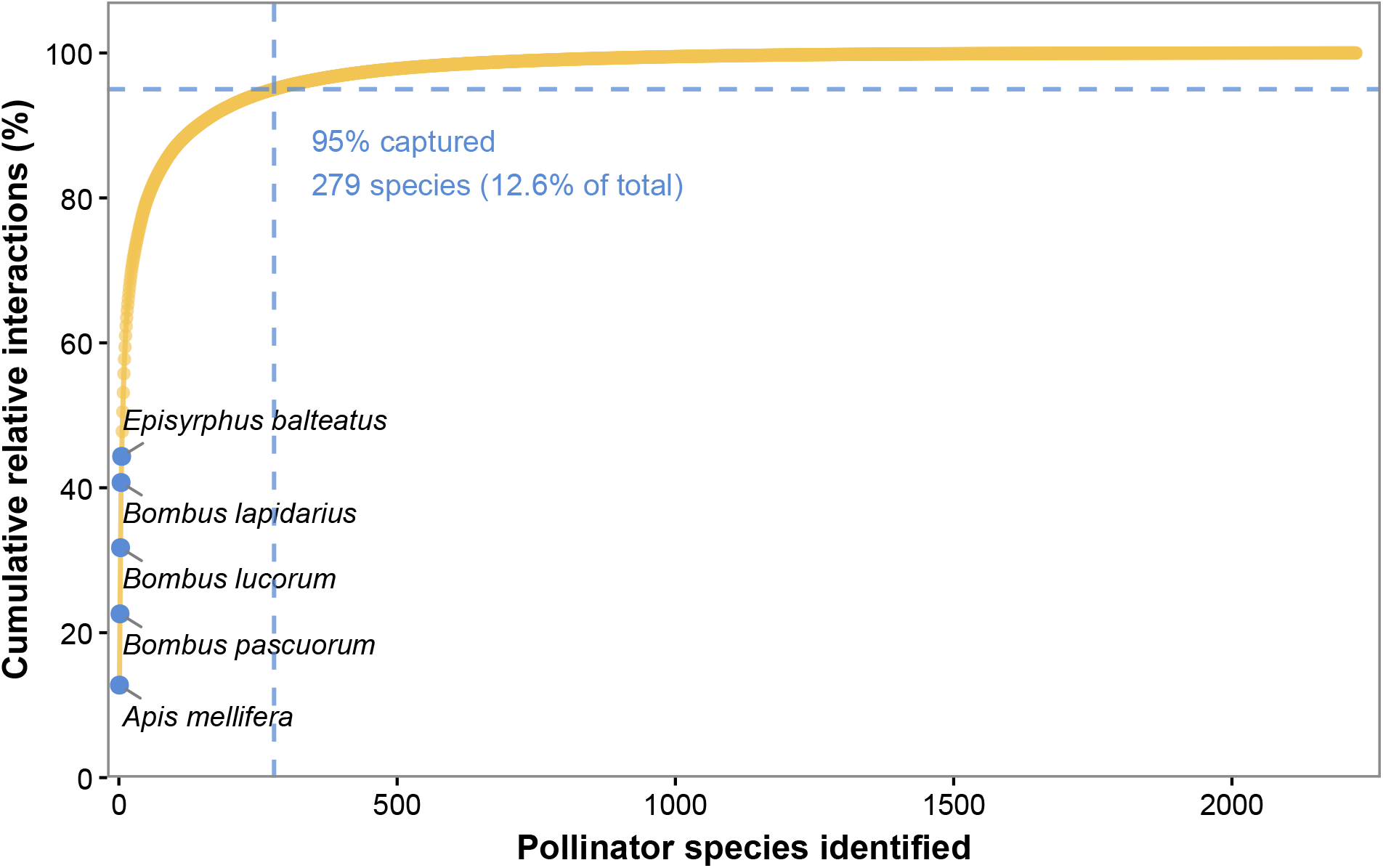
Number of pollinator species required to capture most observed interactions. Pollinator species were ranked according to their mean relative contribution to interactions across all European networks. The cumulative proportion of interactions captured by progressively identifying pollinator species is shown. The dashed lines indicate the threshold of 95% of cumulative interactions, and the highlighted labels indicate the five pollinator species with the highest mean relative interaction contribution.

## Discussion

This study demonstrates that subsampling the plant community, as might be applied for automated pollinator monitoring in the future, is an effective approach for characterizing pollinator richness and the structure of plant-pollinator networks. The best-performing subsampling method was also the simplest; sample the species with highest flowering abundance in the community. Using this approach, sampling interactions on ten species per network was sufficient to capture approximately three quarters of the pollinator richness detected by traditional plant-pollinator network monitoring. Increasing sampling effort further improved the representation of network structure, although gains diminished beyond a certain sampling effort. Overall, sampling the 12-15 most abundant flowering plant species was optimal for capturing pollinator diversity and network metrics across communities with varying size.

The effectiveness of abundance-based sampling at capturing pollinator richness varied greatly across networks (capturing between 23-100% of pollinator richness present in the complete network). Follow-up analyses revealed several reasons why. First, communities that are highly diverse are more under sampled in a monitoring scheme that focuses on sampling a constant number of plant species (i.e., consistently just ten species). Solutions to this issue could involve monitoring designs that sample proportionately to the diversity of the plant community. Second, we identified 47 plant species that are visited by pollinators disproportionately to their abundance, and missing these plant species in the subsample, even if they are rare in the community, will underestimate pollinator diversity. Therefore, automated monitoring practice could benefit from sampling these species when they are present in the community, and incorporating any other prior knowledge of regional plant-pollinator interactions to identify and target plant species that are known to support disproportionately diverse pollinator assemblages. Notably, flower-head species, predominantly from the Asteraceae family, were strongly overrepresented among the rare but highly attractive plants. Asteraceae flower heads are widely recognized as effective pollinator-attraction units, combining conspicuous displays with numerous florets that provide readily accessible floral resources to a broad range of visitors [23,24]. Abundance-based subsampling was also more likely to capture pollinator diversity in highly nested plant-pollinator networks. This pattern is consistent with the organization of nested networks, in which abundant generalist plants are visited by many pollinator species in the community [10,11]. Although nestedness is rarely known *a priori*, historical plant-pollinator network may provide valuable guidance for implementing effective sampling design for automated monitoring.

Contrary to expectations, incorporating flower shape into the subsampling design provided no advantage over abundance-based sampling. This does not imply that floral shape is unimportant for plant-pollinator interactions. Floral shape is known to constrain access to floral resources and thereby filter pollinator assemblages [15,25]. Our results did show the expected significant differences in pollinator community composition across flower-shape categories, and guild-level associations, such as Lepidoptera disproportionately visiting stalk disk flowers and bees disproportionately visiting flag flowers. However, flower shape explained only 2.5% of the variation in pollinator composition among plant species within networks, suggesting that these associations were weak. Pollinator composition varied substantially among plant species within the same flower-shapes category, with “disk flowers open” exhibiting the highest dissimilarity across species within a network. Thus, although flower shape captures some broad ecological differences among pollinator guilds, it is too coarse a predictor of pollinator composition to provide a general advantage for selecting plants for automated monitoring. Additional floral traits, such as flower size, color, or nectar and pollen characteristics, might explain more variation in pollinator assemblages, but incorporating these traits would require substantially more information and effort. For automated monitoring, plant selection criteria must be not only ecologically informative but also practical for field teams. Our results therefore suggest that prioritizing the most abundant flowering plants provides a simple and effective method for plant selection, with little additional benefit from incorporating flower morphology.

Although abundance-based plant selection captured much of the observed pollinator richness, miss rates varied among taxa. Broad representation of a family on the selected plants did not ensure coverage of its constituent species across networks. For example, Apidae contributed the largest number of species-by-network records within Hymenoptera and had a comparatively low pooled miss rate, yet some frequent hymenopterans were missed from a substantial proportion of the networks in which they occurred. These findings suggest that supplementary targeted monitoring may be needed to avoid systematic bias against these species. For instance, several bee species, including social parasites or cleptoparasites (*Bombus bohemicus, B. campestris, B. vestalis, Nomada flavoguttata)*, were frequently missed. The abundance of these species are constrained by the abundance of their hosts, and thus these species may be rare in communities and have low encounter rates [26]. Floral specialists were also frequently missed, such as *Eucera nigrescens*, which show strong associations with Fabaceae [27]. These bees might be missed if their preferred floral hosts are not among the plant species with highest floral abundance. These patterns suggest that automated monitoring focused on abundant plants may under-sample parasitic and specialized taxa, and that targeted supplementary sampling may be needed where such species are conservation priorities.

A relatively small set of pollinator species (279 species; 12.6% of the total) accounted for 95% of the interactions across all networks, offering a concrete target for computer-vision classifier development that would capture the majority of interactions. New image-classification models and approaches are being introduced at a rapid pace. Two key design decisions are whether to constrain predictions to a defined regional species list [28,29] or to use more flexible image-text association models, such as BioCLIP 2 [30], and whether to aim for species-level classification or a coarser taxonomic resolution [31–35]. The concentration of interactions among a relatively small number of pollinator species suggests that AI classifiers focused on a well-curated species list may achieve greater accuracy in species-level identification. However, several *Lasioglossum* species are included in our list of 279 species, yet their identification requires microscopic examination of fine morphological traits [36]. We are therefore cautious about expecting AI classifiers to accurately separate such species in field images, despite their ability to exploit subtle visual cues that are not easily seen by the human eye [37,38].

A potential limitation of automated camera-based monitoring is that it operates at the level of individual flowers or inflorescences rather than at the level of the plant individual or plant population, and visitation can vary substantially among conspecific flowers [39–41], potentially due to multiple factors such as flowering phenology, plant phenotype, and microsite environment [42]. Cameras aimed at just one flower may therefore not represent the overall visitation patterns to that plant species within the community and may also affect inferences about plant-pollinator network structure and stability [43]. Our analyses are based on full interaction networks aggregated at the species level, and as a result, they do not incorporate additional variance arising from individual flowers, and should be interpreted with caution. Further studies are needed to validate the performance of camera-based sampling against traditional field sampling approaches. The complete networks we use as a reference represent the best available approximation of pollinator richness and plant-pollinator interactions for each community, but these reference networks are themselves based on incomplete sampling and may have their own biases [44,45]. For example, transect sampling might disproportionately observe and capture larger, slower or more conspicuous pollinators [46]. Direct comparisons have shown that automated cameras can record more low-frequency interactions than human observers, while still having difficulty detecting small-bodied pollinators [47]. For pollinator richness, our sampling completeness analyses indicate that the reference networks are generally well-sampled, but it is important to recognize that both traditional and automated methods provide filtered views of the true interaction network, and their biases will not be identical.

Looking ahead, a more robust framework for pollinator monitoring is likely to emerge from the integration of traditional and automated sampling approaches within initiatives such as the EU Pollinator Monitoring Scheme (EU PoMS), which aims to establish a harmonized and scalable monitoring system across Europe [48]. Traditional methods remain essential for establishing the regional species pool and obtaining specimens for taxonomic verification, whereas AI-based camera monitoring can extend temporal coverage and resolve fine-scale variation in visitation. Daily sampling periods can influence the composition and structure of plant-pollinator networks [49]. Future studies could therefore test whether the plant species selected for their overall coverage remain representative at different times of day, helping to optimize both plant selection and monitoring schedules. Rather than replacing conventional approaches, automated methods should therefore be viewed as a complementary layer within a multi-method monitoring framework. Systematic evaluations comparing the performance of traditional and automated approaches in sampling pollinators and plant-pollinator networks are still needed. Our results highlight a practical strategy for implementing automated monitoring and identify contexts in which it should be combined with traditional sampling to improve inference on pollinator diversity and interactions.

## Methods

### Data Cleaning and Filtering

EuPPollNet contains 52 studies with 1630 local networks. We grouped the interaction data as the authors of the studies did, typically with different networks representing different sites and years. For our purpose, it was necessary to have an independent assessment on flowering abundance for each plant species. Thus, we retained only the studies of EuPPollNet that conducted a separate survey of floral abundance, which resulted in 36 studies with 1035 networks.

For each network, we identified the most common plant species by averaging flower abundance of each species across all replicate transects within the plant survey. Plant surveys comprised an average of 4.2 replicate transects per network. Studies varied in the methods used to calculate floral abundances (e.g., number of flowers or inflorescences, percent cover of flowers; Supplementary **Table S3**). Although floral abundance was quantified using different metrics across studies, these metrics were used only to estimate relative floral abundance within each network and were not compared among studies. Because scientific names in the plant surveys were not standardized across studies, plant species names were harmonized using the GBIF Backbone Taxonomy through the R package rgbif [50], with unmatched names checked against Plants of the World Online (WFO). Then we checked whether the plant species present in the plant survey data overlapped with those present in the interaction data and excluded networks if more than 25% of the plant species found in the interaction data were absent from the corresponding plant survey. Next, we excluded networks with fewer than 30 interactions from further consideration (Supplementary **Fig. S8**) and those with fewer than ten plant species, which typically resulted from highly limited spatial and temporal grains of sampling (Supplementary **Fig. S9**). We excluded graminoid species because they are predominantly wind-pollinated. All steps in the data cleaning are found in a PRISMA-style flow diagram (Supplementary **Fig. S10**) [51]. The final dataset contained 27 studies with 268 networks distributed across 13 countries (Supplementary **Fig. S11**). Sampling completeness of pollinators was high for these 268 networks (Supplementary **Fig. S12**), suggesting that the sampling intensity of the original networks accurately reflected the diversity of pollinators in their respective subsampled networks.

### Flower shapes

All flowering species were categorized based on their flower shape (taking the accessibility of the reward into account) using the BiolFlor database (www.biolflor.de) [52], accessed through the R package “TR8” [53]. The classification follows the flower-shape scheme of Kugler’s methodology [54,55]. Accordingly, flower shapes were classified into 10 categories (Supplementary **Table S4**). Missing flower shape information was manually assigned based on closely related congeneric taxa, which typically share similar flower shape. Networks contained an average of 7.5 flower-shape categories (Supplementary **Fig. S13**). Across all networks, the most common flower-shape categories were *flower head, flag flowers, and disk flower hidden*, which were primarily represented by species from the families Asteraceae, Fabaceae, and Rosaceae (supplementary **Fig. S13**).

### Pollinator richness across sampling strategies

Using the complete plant-pollinator interaction networks reported in the original studies, we evaluated three ecologically informed sampling strategies as proxies for pollinator communities. Each strategy was evaluated across three sampling effort levels (three, five and ten plant species) and compared their performance with a random-selection benchmark. Specifically, we tested: (i) abundance-based, selecting the most abundant flowering plant species; (ii) flower-shape abundance-based, selecting abundant plant species representing different flower shapes; and (iii) phylogenetic distance based, selecting the most phylogenetically distant plant species, as it may help as a surrogate of unmeasured reproductive traits influencing pollinator interactions. To provide a benchmark for comparison, we also randomly selected three, five and ten plant species. For each sampling effort, the procedure was repeated 1000 times, and the percentage of the total pollinator species richness represented by the selected plants was quantified for each iteration and averaged across iterations. Finally, complete networks were used as the reference for quantifying the proportion of total pollinator species richness captured by each sampling strategy.

We used two-sided paired Wilcoxon signed-rank tests to compare the proportion of total pollinator richness captured across networks by the different sampling strategies. Comparisons were paired by network, and only networks with non-missing values were included. For the sampling effort of three and five plant species, we conducted all pairwise comparisons among sampling strategies but for ten plant species, comparisons were restricted to the abundance-based and random sampling, as too few networks contained ten or more distinct flower shape categories.

### Network structure across sampling strategies

We used three widely used and complementary network-level indices to assess plant-pollinator interaction network structure across sampling strategies: connectance, nestedness and selectivity. Connectance measures the proportion of realized links in a network [56] and it is highly sensitive to sampling completeness, providing a sensitive measure of sampling performance [57,58]. Nestedness, quantified as NODF metric [59], describes the tendency of specialist species to interact with subsets of the partners utilized by more generalized species [60–62], which is comparatively robust to sampling incompleteness [58,63]. Selectivity was quantified using H , which quantifies the degree to which species interact selectively with available partners based on interaction frequencies and is largely independent of network size, providing a robust measure of overall network specialization [64]. Together, these indices capture complementary dimensions of network structure and allow us to assess whether different subsampling strategies preserve network properties that vary in their sensitivity to sampling effort. We calculated all three metrics at the network level using the bipartite R package [65] and compared values from subsampled networks with those from complete networks. We quantified the strength of the association with Spearman’s rank correlation to assess how well subsampling preserved the relative ranking of networks despite differences in absolute metric values.

To evaluate how sampling effort influenced the representation of network structure, we performed a progressive abundance-based selection analysis. For each network, plant species were ranked by scaled flower abundance, and the top n most abundant plant species were selected to construct a sub-network, with *n* increasing sequentially from 1 to all plant species in the network (or maximum of 50 species). For each value of *n*, we reconstructed subnetworks by selecting either the *n* most abundant plant species or *n* plant species at random. We calculated connectance, nestedness (NODF), and network-level specialization (H_2_) for each subnetwork and calculated Spearman’s rank correlation coefficients () between subnetwork and complete-network values for each metric across all eligible networks. For each selection strategy and metric, we identified the smallest observed number of selected plant species at which reached or exceeded 0.8, which we considered a strong association.

### Factors associated with plant sampling effectiveness

We conducted a set of follow-up analyses focusing on the best-performing strategy (i.e., abundance-based sampling) to explore how community characteristics are associated with its effectiveness. We first tested how subsampled pollinator richness varied with the number of plant and pollinator species in the complete network and with network nestedness. Associations in both analyses were quantified using Pearson correlation coefficients. Then, we assessed how subsampled pollinator richness varied with the presence of rare but highly attractive plants (i.e., plants receiving disproportionately many interactions relative to their abundance). We first fitted a linear mixed-effects model with subsampled pollinator richness as response variable and floral abundance as fixed effect. To account for strong right-skewness in floral abundance, we transformed floral abundance using a log (x + 1) transformation followed by z-score standardization. Study identity and network identity (site × year) nested within study were included as random intercepts to account for the hierarchical structure and non-independence of observations. Residuals were calculated as the difference between observed and model-predicted values, with positive residuals indicating plant species attracting higher-than-expected pollinator richness given their abundance. “Rare-Attractive” plant species were identified as those with (i) standardized floral abundance below the 5th percentile across all observations and (ii) residuals above the 90th percentile (Supplementary **Fig. S14**). A total of 47 plant species met the “rare-attractive” criteria (Supplementary **Table S5**). Finally, we tested whether omission of “rare-attractive” plants affected the percentage of pollinators captured. Following this classification, we divided the subsampled networks into two groups: those that missed at least one “rare-attractive” plant and those that did not, either because no “rare-attractive” plants were present in the original network or because the sampling strategy captured all of them. Differences between the two groups were tested using Wilcoxon rank-sum tests. Analogous analyses with varying percentile ranges yielded qualitatively similar results (Supplementary **Fig. S14**).

Finally, we tested if overlap in pollinator assemblages across flower shapes could explain the limited improvement of incorporating flower-shape into abundance-based sampling. Pollinator assemblages were characterized at the genus level using presence-absence data for each plant species within each study network. Thus, each observation represented a plant species × network combination. Jaccard dissimilarities were analyzed using PERMANOVA with 999 permutations. Study identity was included as a fixed covariate, and permutations were restricted within networks. Pairwise comparisons used the same design with Benjamini-Hochberg adjusted P values. We also tested for homogeneity of multivariate dispersion using *betadisper*, with permutations similarly restricted within networks, to assess whether any PERMANOVA effect could be confounded by differences in within-group dispersion [66]. To further determine whether plants sharing the same flower shape attracted similar pollinator assemblages within networks, we calculated mean pairwise Sørensen dissimilarity for each network × flower-shape combination. Differences among flower shapes were tested using a linear mixed-effects model, with flower shape as a fixed effect and study and network identities as random intercepts. Pairwise estimated marginal means were compared using Sidak-adjusted P values.

### Target pollinator species for AI classification

We examined families in the major pollinator orders that were consistently missed when sampling the ten most abundant plant species, with particular attention to Apidae because this ecologically important and diverse bee family accounted for most of the interactions in the dataset (71.4%). For each species, we calculated a miss rate as the proportion of networks in which it occurred under complete sampling but was not captured by the sampling strategy. Miss rates were then summarized at the family level to identify families that were consistently underrepresented.

To identify pollinator species that should be prioritized for AI classifier development, we quantified the continental-scale contribution of each pollinator species to recorded plant-pollinator interactions across Europe. From the 1,630 networks in EuPPollNet database, we calculated the relative interaction contribution of each pollinator species as its number of recorded interactions divided by the total number of interactions in that network. Species absent from a network were assigned a relative contribution of zero. We then calculated the mean relative interaction contribution of each species across networks, ranked species in descending order of this value, and constructed an accumulation contribution curve. Pollinator species were retained and reported until their cumulative mean relative contribution accounted for 95% of the total mean relative contribution.

## Acknowledgments

We thank all contributors to the EuPPolNet database for providing the data used in this study. All flower and insect icons used in the main-text figures were hand-drawn by Y.Z. This research was supported by Biodiversa+ through the SEPPI project (Standardised European monitoring of plant-pollinator interactions), co-funded by the European Commission and the Deutsche Forschungsgemeinschaft (DFG, project 532239370). Y.Z. was additionally supported by a scholarship from the China Scholarship Council (CSC). J.B.L. was additionally supported by a Juan de la Cierva fellowship funded by the Spanish Ministry of Science.

## Author contributions

Y.Z. contributed to the conceptualization of the study, conducted the formal analyses, prepared the figures, and wrote the first draft of the manuscript. J.B.L. contributed to the study design, analytical approach, code development, figure presentation, and manuscript revision. J.M.H. contributed to the analytical framework and manuscript writing. W.G. and D.R. contributed to data analysis and manuscript revision. T.M.K. conceptualized and supervised the study, guided the overall analytical framework, and critically revised the manuscript. All authors reviewed and approved the final manuscript.

## Data availability

The plant-pollinator interaction data analyzed in this study were obtained from the EuPPollNet database and are available on Zenodo (https://doi.org/10.5281/zenodo.15183272).

## Code availability

All R scripts used for data processing, statistical analyses, and figure generation are publicly available at https://github.com/YupingZhong/AIPlantSelection. The final version of the code will be deposited in Zenodo and made publicly available upon publication.

## References

1. van Klink, R. et al. Emerging technologies revolutionise insect ecology and monitoring. Trends Ecol. Evol. 37, 872–885 (2022).

2. Ştefan, V., Workman, A., Cobain, J. C., Rakosy, D. & Knight, T. M. Utilising affordable smartphones and open-source time-lapse photography for pollinator image collection and annotation. J. Pollinat. Ecol. 38, 1–21 (2025).

3. Steen, R. Diel activity, frequency and visit duration of pollinators in focal plants: in situ automatic camera monitoring and data processing. Methods Ecol. Evol. 8, 203–213 (2017).

4. Lortie, C. J., Budden, A. & Reid, A. From birds to bees: applying video observation techniques to invertebrate pollinators. J. Pollinat. Ecol. 6, 125–128 (2012).

5. Stark, T. et al. Utilizing CNNs for classification and uncertainty quantification for 15 families of European fly pollinators. PLOS ONE 20, e0323984 (2025).

6. Bjerge, K., Karstoft, H., Mann, H. M. R. & Høye, T. T. A deep learning pipeline for time-lapse camera monitoring of insects and their floral environments. Ecol. Inform. 84, 102861 (2024).

7. Peralta, G. et al. Trait matching and phenological overlap increase the spatio-temporal stability and functionality of plant–pollinator interactions. Ecol. Lett. 23, 1107–1116 (2020).

8. Lázaro, A., Gómez-Martínez, C., Alomar, D., González-Estévez, M. A. & Traveset, A. Linking species-level network metrics to flower traits and plant fitness. J. Ecol. 108, 1287–1298 (2020).

9. Olito, C. & Fox, J. W. Species traits and abundances predict metrics of plant–pollinator network structure, but not pairwise interactions. Oikos 124, 428–436 (2015).

10. Krishna, A., Guimarães Jr, P. R., Jordano, P. & Bascompte, J. A neutral-niche theory of nestedness in mutualistic networks. Oikos 117, 1609–1618 (2008).

11. Vázquez, D. P. et al. Species abundance and asymmetric interaction strength in ecological networks. Oikos 116, 1120–1127 (2007).

12. Wang, H. et al. Complex floral traits shape pollinator attraction to flowering plants in urban greenspaces. Urban For. Urban Green. 91, 128165 (2024).

13. Howard, S. R. et al. Honeybees prefer novel insect-pollinated flower shapes over bird-pollinated flower shapes. Curr. Zool. 65, 457–465 (2019).

14. Lehrer, M., Horridge, G. A., Zhang, S. W. & Gadagkar, R. Shape vision in bees: innate preference for flower-like patterns. Philos. Trans. R. Soc. Lond. B. Biol. Sci. 347, 123–137 (1995).

15. Krishna, S. & Keasar, T. Morphological Complexity as a Floral Signal: From Perception by Insect Pollinators to Co-Evolutionary Implications. Int. J. Mol. Sci. 19, 1681 (2018).

16. Fenster, C. B., Armbruster, W. S., Wilson, P., Dudash, M. R. & Thomson, J. D. Pollination Syndromes and Floral Specialization. Annu. Rev. Ecol. Evol. Syst. 35, 375–403 (2004).

17. Srivastava, D. S., Cadotte, M. W., MacDonald, A. A. M., Marushia, R. G. & Mirotchnick, N. Phylogenetic diversity and the functioning of ecosystems. Ecol. Lett. 15, 637–648 (2012).

18. Mollenhauer, H. et al. Long-term environmental monitoring infrastructures in Europe: observations, measurements, scales, and socio-ecological representativeness. Sci. Total Environ. 624, 968–978 (2018).

19. Lanuza, J. B. et al. EuPPollNet: A European Database of Plant-Pollinator Networks. Glob. Ecol. Biogeogr. 34, e70000 (2025).

20. Ward, D. F. Understanding sampling and taxonomic biases recorded by citizen scientists. J. Insect Conserv. 18, 753–756 (2014).

21. Boakes, E. H. et al. Patterns of contribution to citizen science biodiversity projects increase understanding of volunteers’ recording behaviour. Sci. Rep. 6, 33051 (2016).

22. Van Horn, G. et al. The iNaturalist Species Classification and Detection Dataset. in 8769–8778 (IEEE, 2018). doi:10.1109/cvpr.2018.00914.

23. Elomaa, P., Zhao, Y. & Zhang, T. Flower heads in Asteraceae—recruitment of conserved developmental regulators to control the flower-like inflorescence architecture. Hortic. Res. 5, 36 (2018).

24. Fu, L. et al. Let’s pluck the daisy: dissection as a tool to explore the diversity of Asteraceae capitula. Bot. J. Linn. Soc. 201, 391–399 (2023).

25. Stang, M., Klinkhamer, P. G. L. & Van Der Meijden, E. Size constraints and flower abundance determine the number of interactions in a plant–flower visitor web. Oikos 112, 111–121 (2006).

26. Sheffield, C. S., Pindar, A., Packer, L. & Kevan, P. G. The potential of cleptoparasitic bees as indicator taxa for assessing bee communities. Apidologie 44, 501–510 (2013).

27. Dorchin, A. et al. Bee flowers drive macroevolutionary diversification in long-horned bees. Proc. R. Soc. B Biol. Sci. 288, 20210533 (2021).

28. Mac Aodha, O., Cole, E. & Perona, P. Presence-Only Geographical Priors for Fine-Grained Image Classification. in 9595–9605 (IEEE, 2019). doi:10.1109/iccv.2019.00969.

29. Sun, J., Futahashi, R. & Yamanaka, T. Improving the Accuracy of Species Identification by Combining Deep Learning With Field Occurrence Records. Front. Ecol. Evol. 9, 762173 (2021).

30. Gu, J. et al. BioCLIP 2: Emergent Properties from Scaling Hierarchical Contrastive Learning. in vol. 38 102778–102811 (Curran Associates, Inc., 2025).

31. Buschbacher, K., Ahrens, D., Espeland, M. & Steinhage, V. Image-based species identification of wild bees using convolutional neural networks. Ecol. Inform. 55, 101017 (2020).

32. Spiesman, B. J. et al. Assessing the potential for deep learning and computer vision to identify bumble bee species from images. Sci. Rep. 11, 7580 (2021).

33. Stark, T. et al. YOLO object detection models can locate and classify broad groups of flower-visiting arthropods in images. Sci. Rep. 13, 16364 (2023).

34. Shirali, H. et al. Image-based recognition of parasitoid wasps using advanced neural networks. Invertebr. Syst. 38, IS24011 (2024).

35. Bjerge, K. et al. Hierarchical classification of insects with multitask learning and anomaly detection. Ecol. Inform. 77, 102278 (2023).

36. Michener, C. D. The Bees of the World. (Johns Hopkins University Press, Baltimore, 2007). doi:10.56021/9780801885730.

37. Valan, M., Makonyi, K., Maki, A., Vondráček, D. & Ronquist, F. Automated Taxonomic Identification of Insects with Expert-Level Accuracy Using Effective Feature Transfer from Convolutional Networks. Syst. Biol. 68, 876–895 (2019).

38. Geirhos, R. et al. ImageNet-trained CNNs are biased towards texture; increasing shape bias improves accuracy and robustness. in International conference on learning representations (2019).

39. Harder, L. D. & Barrett, S. C. H. Mating cost of large floral displays in hermaphrodite plants. Nature 373, 512–515 (1995).

40. Kunin, W. E. Population Size and Density Effects in Pollination: Pollinator Foraging and Plant Reproductive Success in Experimental Arrays of Brassica Kaber. J. Ecol. 85, 225–234 (1997).

41. Lázaro, A., Lundgren, R. & Totland, ø. Co-flowering neighbors influence the diversity and identity of pollinator groups visiting plant species. Oikos 118, 691–702 (2009).

42. Arroyo-Correa, B., Bartomeus, I. & Jordano, P. Individual-based plant–pollinator networks are structured by phenotypic and microsite plant traits. J. Ecol. 109, 2832–2844 (2021).

43. Arroyo-Correa, B., Jordano, P. & Bartomeus, I. Intraspecific variation in species interactions promotes the feasibility of mutualistic assemblages. Ecol. Lett. 26, 448–459 (2023).

44. Westphal, C. et al. Measuring bee diversity in different european habitats and biogeographical regions. Ecol. Monogr. 78, 653–671 (2008).

45. Olesen, J. M., Bascompte, J., Elberling, H. & Jordano, P. Temporal Dynamics in a Pollination Network. Ecology 89, 1573–1582 (2008).

46. Thompson, A. et al. Pollinator sampling methods influence community patterns assessments by capturing species with different traits and at different abundances. Ecol. Indic. 132, 108284 (2021).

47. Serra-Marin, P. E. et al. Comparative assessment of automated and manual monitoring in comprehensive plant–pollinator communities. Methods Ecol. Evol. 16, 2960–2978 (2025).

48. Potts, S. G. et al. Refined Proposal for an EU Pollinator Monitoring Scheme. https://publications.jrc.ec.europa.eu/repository/handle/JRC138660 (2024) doi:10.2760/2005545.

49. Ballarin, C. S. et al. Optimising daily sampling of plant–bee interaction networks. J. Anim. Ecol. https://doi.org/10.1111/1365-2656.70309 (2026) doi:10.1111/1365-2656.70309.

50. Chamberlain, S., Oldoni, D. & Waller, J. rgbif: Interface to the Global Biodiversity Information Facility API. CRAN Contrib. Packag. https://doi.org/10.32614/cran.package.rgbif (2012) doi: 10.32614/cran.package.rgbif.

51. Page, M. J. et al. The PRISMA 2020 statement: an updated guideline for reporting systematic reviews. BMJ 372, 71 (2021).

52. Kühn, I., Durka, W. & Klotz, S. BiolFlor – a new plant-trait database as a tool for plant invasion ecology. Divers. Distrib. 10, 363–365 (2004).

53. Bocci, G. TR8: an R package for easily retrieving plant species traits. Methods Ecol. Evol. 6, 347–350 (2015).

54. Kugler, H. Einführung in die Blütenökologie. (Gustav Fischer, Stuttgart, 1955).

55. Kugler, H. Blütenökologie. (Gustav Fischer, Stuttgart, 1970).

56. Dunne, J. A., Williams, R. J. & Martinez, N. D. Food-web structure and network theory: The role of connectance and size. Proc. Natl. Acad. Sci. U. S. A. 99, 12917–12922 (2002).

57. Blüthgen, N., Fründ, J., Vázquez, D. P. & Menzel, F. What do interaction network metrics tell us about specialization and biological traits? Ecology 89, 3387–3399 (2008).

58. Rivera-Hutinel, A., Bustamante, R. O., Marín, V. H. & Medel, R. Effects of sampling completeness on the structure of plant–pollinator networks. Ecology 93, 1593–1603 (2012).

59. Almeida-Neto, M., Guimarães, P., Guimarães, P. R., Loyola, R. D. & Ulrich, W. A consistent metric for nestedness analysis in ecological systems: reconciling concept and measurement. Oikos 117, 1227–1239 (2008).

60. Bascompte, J., Jordano, P., Melián, C. J. & Olesen, J. M. The nested assembly of plant–animal mutualistic networks. Proc. Natl. Acad. Sci. 100, 9383–9387 (2003).

61. Ollerton, J. The Pollination Ecology of an Assemblage of Grassland Asclepiads in South Africa. Ann. Bot. 92, 807–834 (2003).

62. Jordano, P., Bascompte, J. & Olesen, J. M. The ecological consequences of complex topology and nested structure in pollination webs. in Plant–Pollinator Interactions: From Specialization to Generalization (eds Waser, N. M. & Ollerton, J.) 173–199 (University of Chicago Press, Chicago, 2006).

63. Nielsen, A. & Bascompte, J. Ecological networks, nestedness and sampling effort. J. Ecol. 95, 1134–1141 (2007).

64. Blüthgen, N., Menzel, F. & Blüthgen, N. Measuring specialization in species interaction networks. BMC Ecol. 6, 9 (2006).

65. Dormann, C. F., Gruber, B. & Fründ, J. Introducing the bipartite package: Analysing ecological networks. R News 8, 8–11 (2008).

66. Anderson, M. J. Distance-Based Tests for Homogeneity of Multivariate Dispersions. Biometrics 62, 245–253 (2006).

