## Supplementary Figures and Tables for "Optimizing plant species selection for automated monitoring of plant-pollinator interactions"

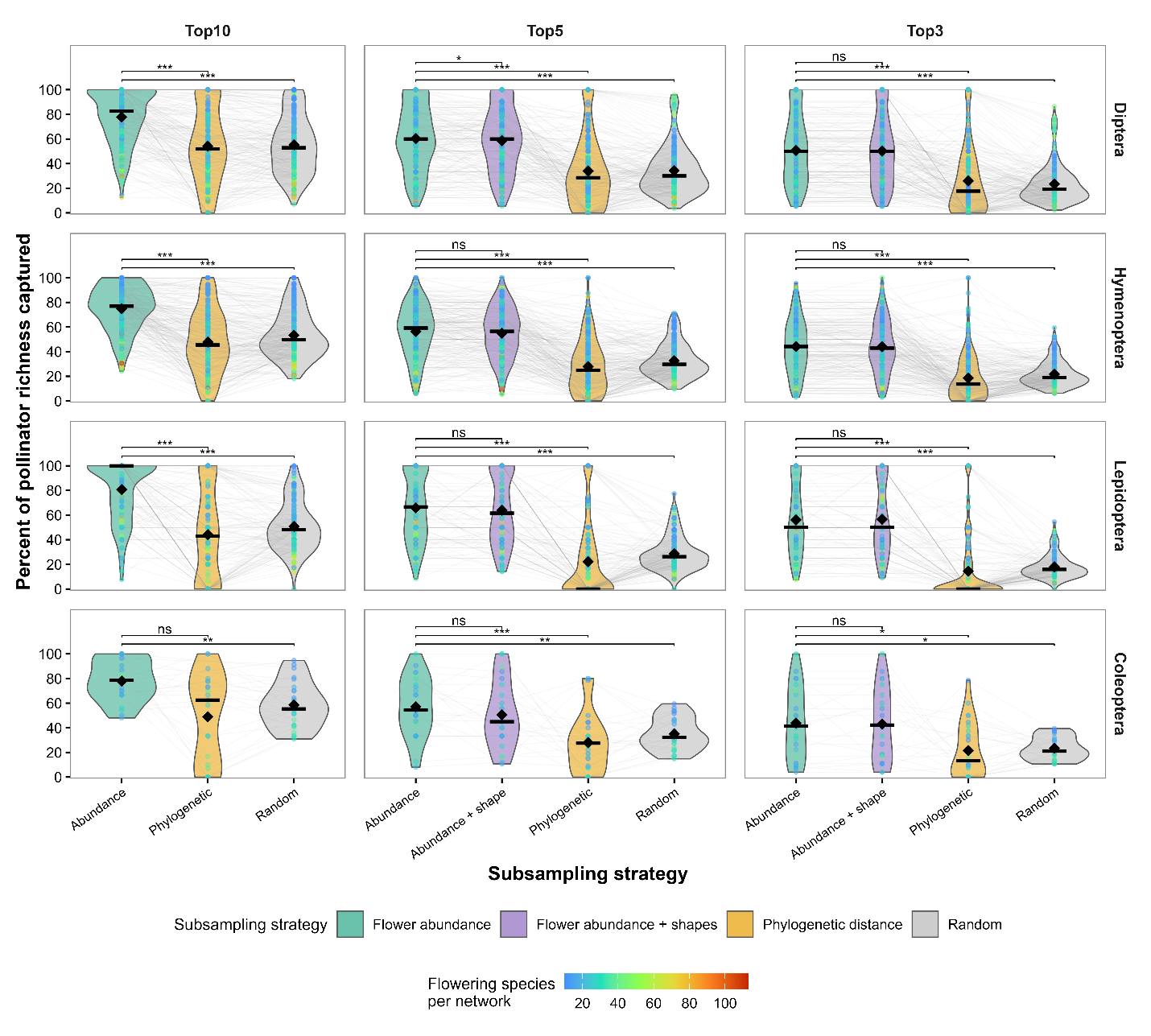

**Supplementary Figure S1.** **Percentage of pollinator species richness captured under alternative plant-selection strategies, shown separately for four insect orders. Rows represent Diptera, Hymenoptera, Lepidoptera, and Coleoptera, and columns represent selection of ten, five, and three plant species.** Violin colors indicate abundance-based selection (green), selection combining floral abundance and flower shape (purple), phylogenetic distance-based selection (yellow), and random selection (gray). Points represent individual networks, with colors indicating flowering plant species richness in the floral-survey data. Gray lines connect results from the same network across strategies. Black diamonds and horizontal bars indicate means and medians, respectively. Brackets indicate selected pairwise comparisons conducted using two-sided paired Wilcoxon signed-rank tests with a normal approximation and continuity correction. Each comparison included only networks with non-missing values for both strategies. P values were not adjusted for multiple comparisons. Significance labels indicate P < 0.001 (***), 0.001 ≤ P < 0.01 (**), 0.01 ≤ P < 0.05 (*), and P ≥ 0.05 (ns). The combined abundance-and-flower-shape strategy is not shown for selection of 10 plant species because too few networks contained ten or more distinct flower-shape categories.

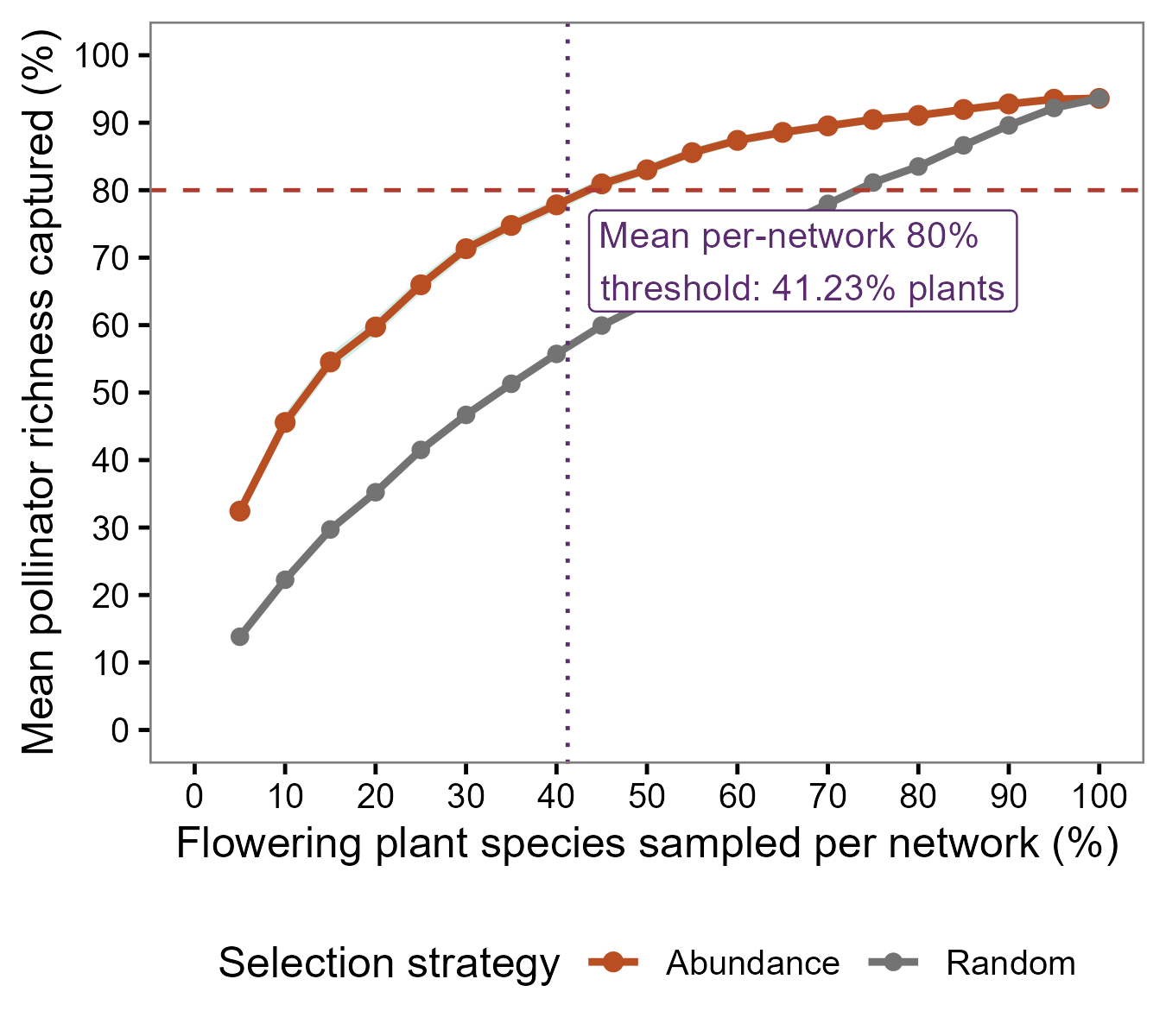

**Supplementary Figure S2. Pollinator richness captured under proportional plant subsampling.** The percentage of pollinator species captured was evaluated across plant–pollinator networks at sampling proportions ranging from 5% to 100% of the flowering plant community. Plants were selected either in descending order of local floral abundance (orange) or randomly (gray). Points and lines show the mean percentage of pollinator richness captured across networks at each sampling proportion; random-selection values were averaged across 50 independent repetitions. The horizontal dashed line indicates the target of 80% pollinator richness. For each network, we identified the minimum proportion of plants required for abundance-based sampling to reach this target. The vertical dotted line indicates the mean of these network-specific thresholds (41.23%).

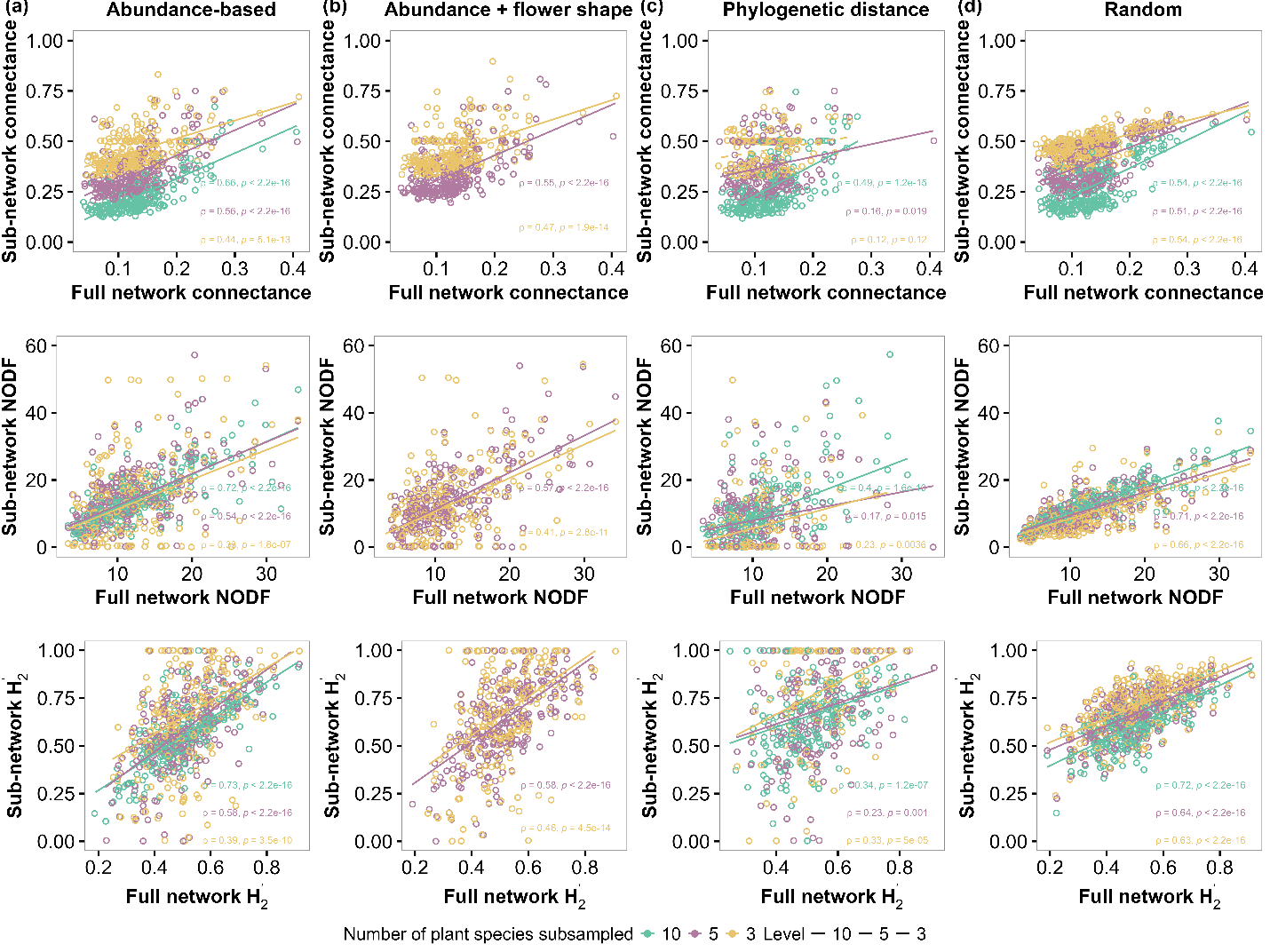
 **Supplementary Figure S3. Associations between subnetwork and full-network structural descriptors under different plant-selection strategies and sampling efforts.** Subnetwork descriptors are plotted against the corresponding full-network values for connectance, nestedness (NODF) and network-level specialization (H2′) under four plant-selection strategies: (a) selection based on floral abundance, (b) selection based on floral abundance across flower-shape categories, (c) selection maximizing phylogenetic distances among plants and (d) random selection. Each point represents one network, and colors indicate the number of plant species sampled. Spearman’s rank correlation coefficients (ρ) and corresponding *P* values are reported separately for each sampling effort. Lines show linear regression fits for visualization only and were not used to calculate the reported correlations.

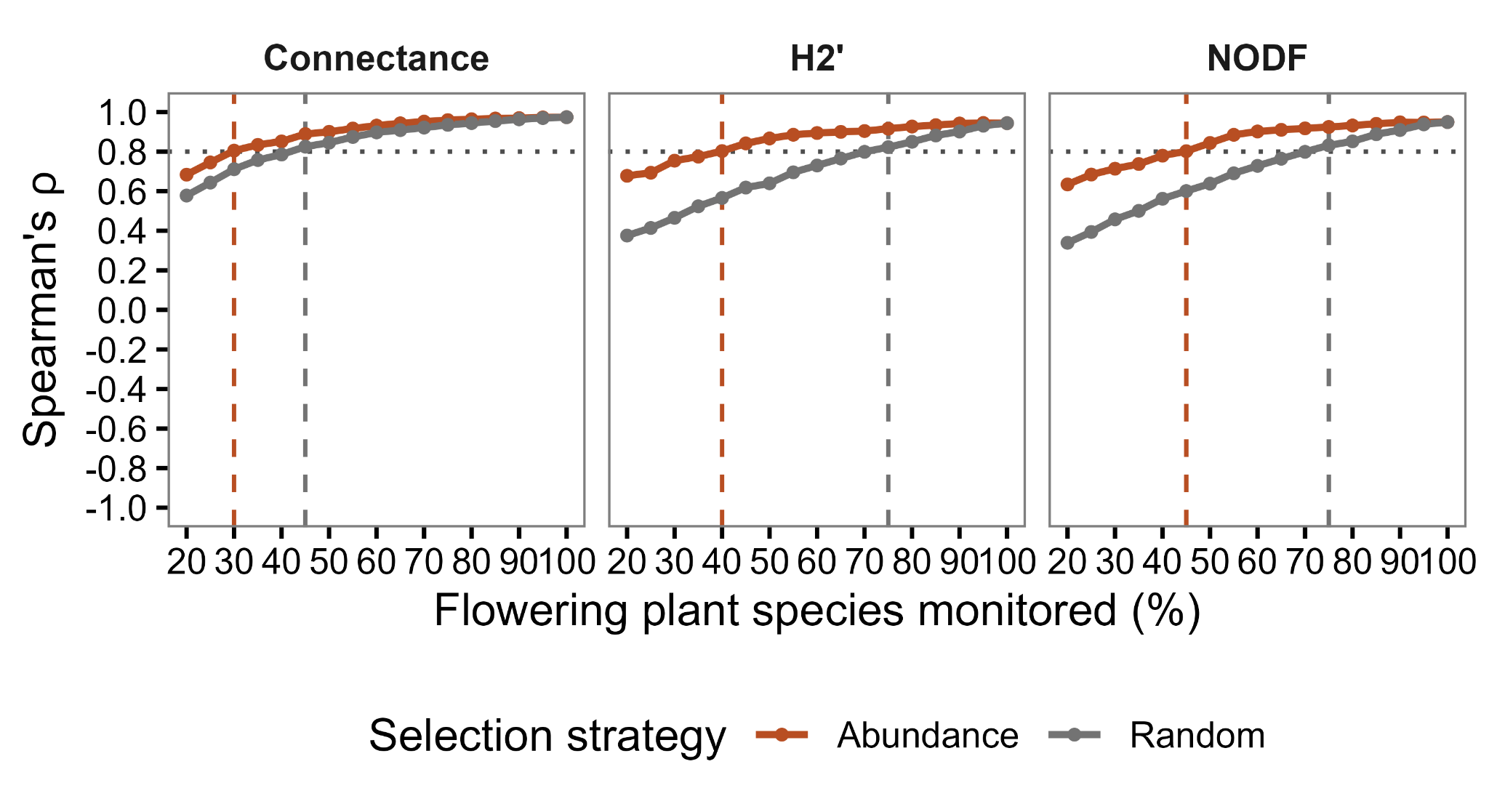

**Supplementary Figure S4. Correspondence between subnetwork and full-network structural descriptors across proportional plant-sampling efforts.** Spearman’s rank correlations (ρ) between descriptors calculated from subsampled and complete plant–pollinator networks are shown for connectance, network-level specialization (H2′) and nestedness (NODF). Flowering plants were selected either in descending order of local floral abundance (orange) or randomly (gray). Points and lines show correlations across networks at each sampling proportion; values for random selection represent means across 50 independent sampling repetitions. The horizontal dotted line indicates the threshold for strong rank-order correspondence (ρ = 0.8). Vertical dashed lines indicate the minimum observed sampling proportion at which each strategy reached this threshold for the corresponding network descriptor.

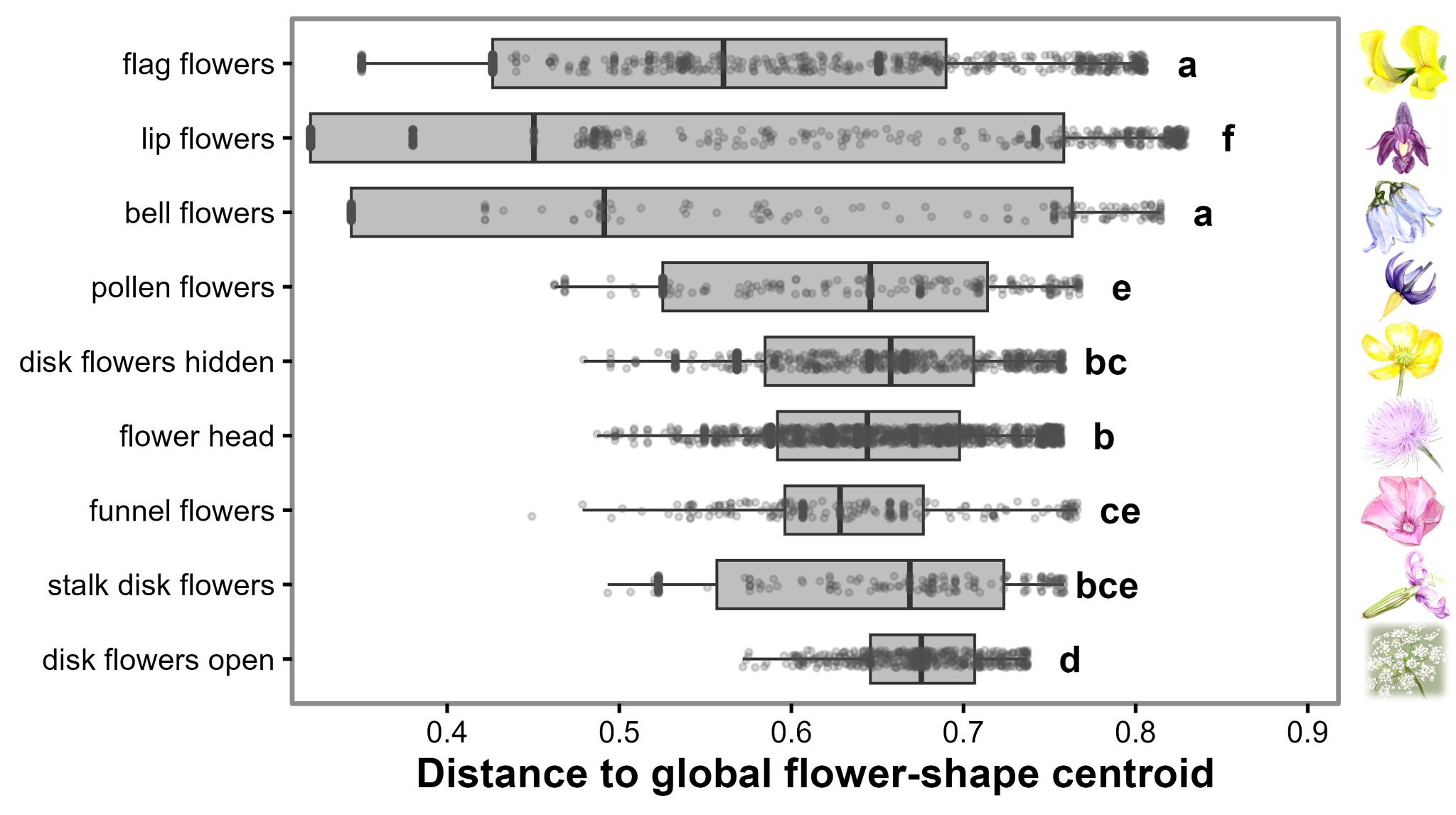

**Supplementary Figure S5. Multivariate dispersion of pollinator assemblage composition among flower-shape categories.** Multivariate dispersion was quantified as the bias-adjusted Jaccard distance of each plant species × network observation to the global centroid of its flower-shape category. Boxplots show the median and interquartile range, whiskers extend to 1.5 times the interquartile range, and points represent individual plant species × network observations. Dispersion differed significantly among flower-shape categories (PERMDISP: *F* = 125.77, *P* = 0.001; 999 permutations restricted within study networks). Letters indicate pairwise PERMDISP groupings based on Benjamini–Hochberg-adjusted permutation *P* values; categories sharing at least one letter did not differ significantly. Letters represent statistical groupings rather than an ordered ranking of dispersion. This analysis was used as a diagnostic for the PERMANOVA and does not directly quantify dissimilarity among plant species within individual networks.

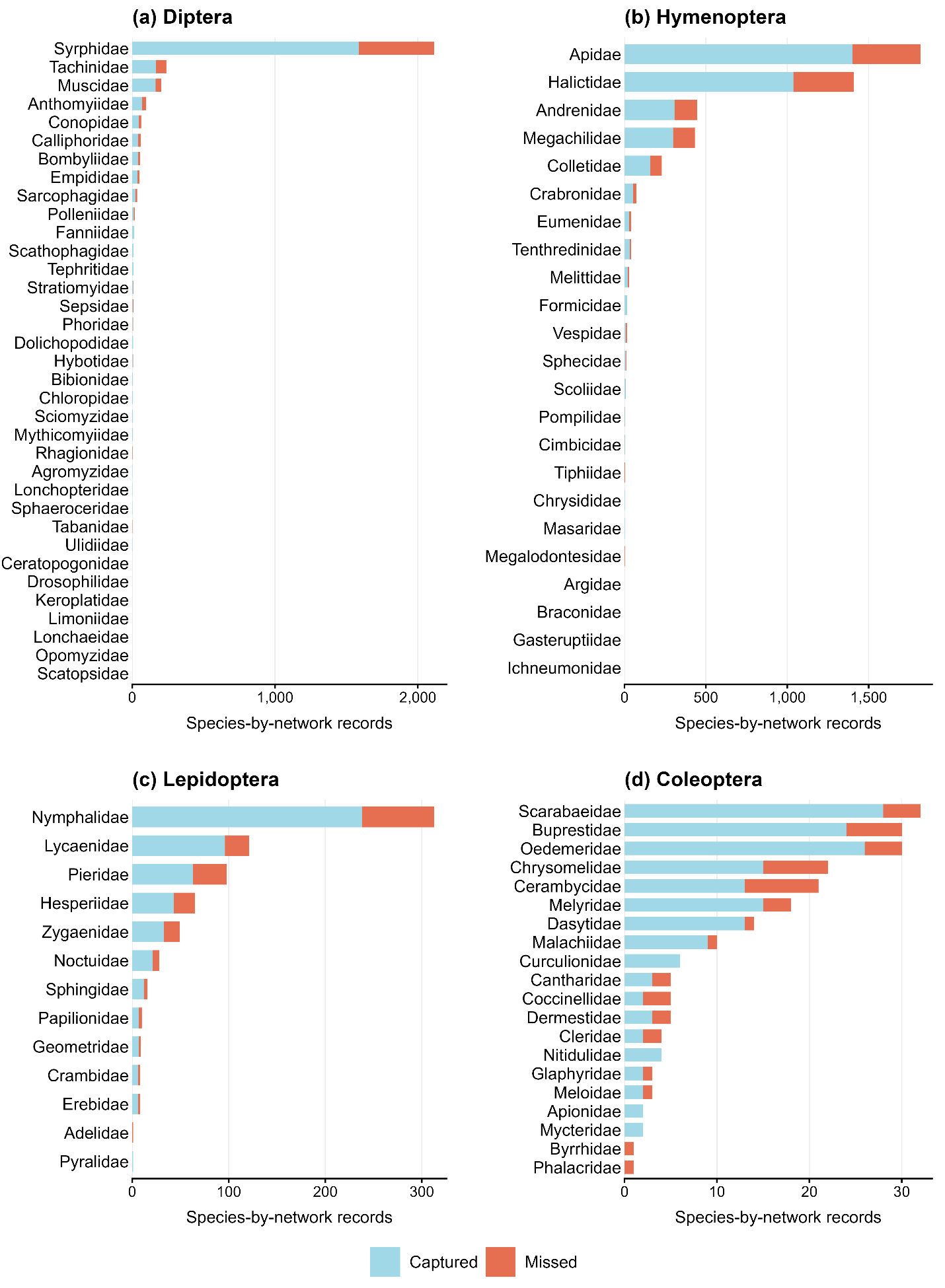

**Figure S6. Family-level representation of insect pollinators when monitoring the ten most abundant plant species.** Each pollinator species recorded in a network was treated as a separate species-by-network record. A record was classified as captured if the species had at least one recorded interaction with the ten selected plant species in that network, and as missed otherwise. Bars show the numbers of captured (blue) and missed (orange) records, grouped by family within (a) Diptera, (b) Hymenoptera, (c) Lepidoptera, and (d) Coleoptera. Count axes differ among panels to accommodate differences in record numbers. Families are ordered by decreasing total record count within each order. Families with zero missed records are retained.

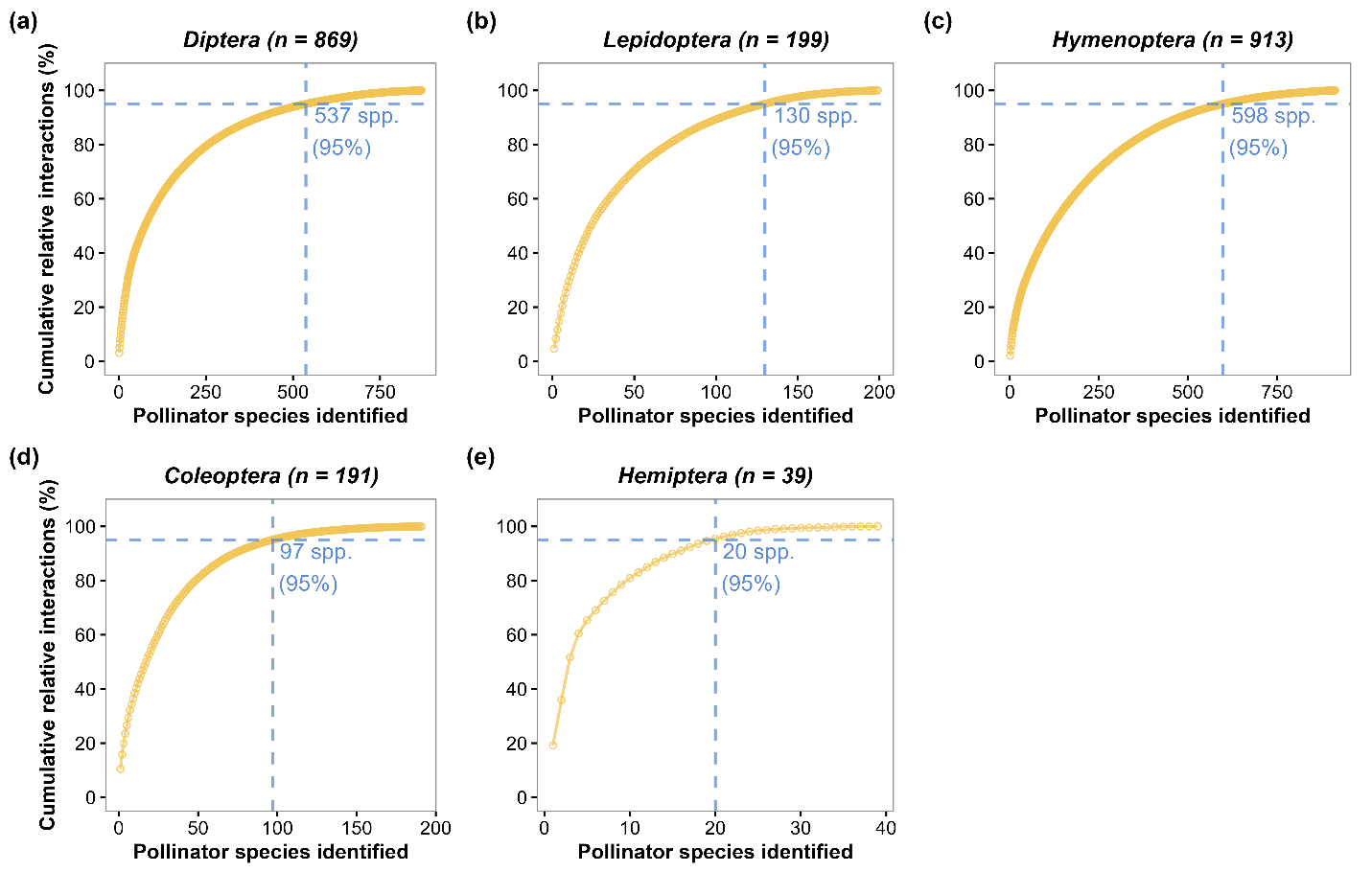

**Supplementary Figure S7.** **Cumulative contribution of pollinator species to mean relative interactions, shown separately for Diptera, Lepidoptera, Hymenoptera, Coleoptera, and Hemiptera.** Species were ranked by their mean relative interaction contribution within each order. Vertical and horizontal dashed lines indicate the number of species required to account for 95% of interactions. Panels represent the five most frequently recorded pollinator orders in the dataset: (a) Diptera (flies), (b) Lepidoptera (butterflies and moths), (c) Hymenoptera (bees and wasps), (d) Coleoptera (beetles), and (e) Hemiptera (true bugs).

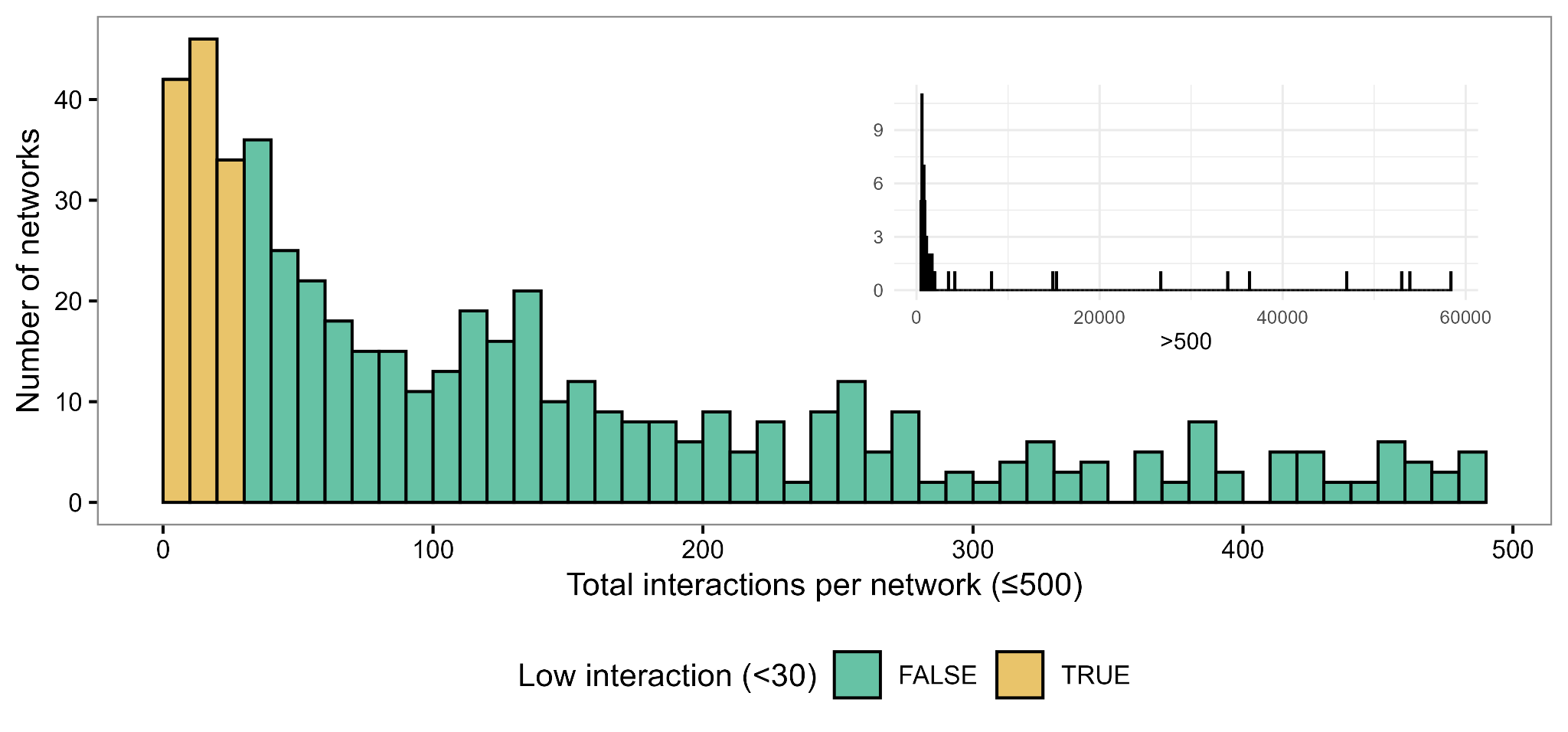

**Supplementary Figure S8.** Distribution of the total number of recorded interactions per network after matching the interaction and floral-abundance datasets and excluding networks in which more than 25% of plant taxa recorded in the interaction data were absent from the floral survey. Networks with fewer than 30 interactions were excluded, whereas those with at least 30 interactions were retained for subsequent filtering. The main panel shows networks with up to 500 interactions, and the inset shows networks with more than 500 interactions.

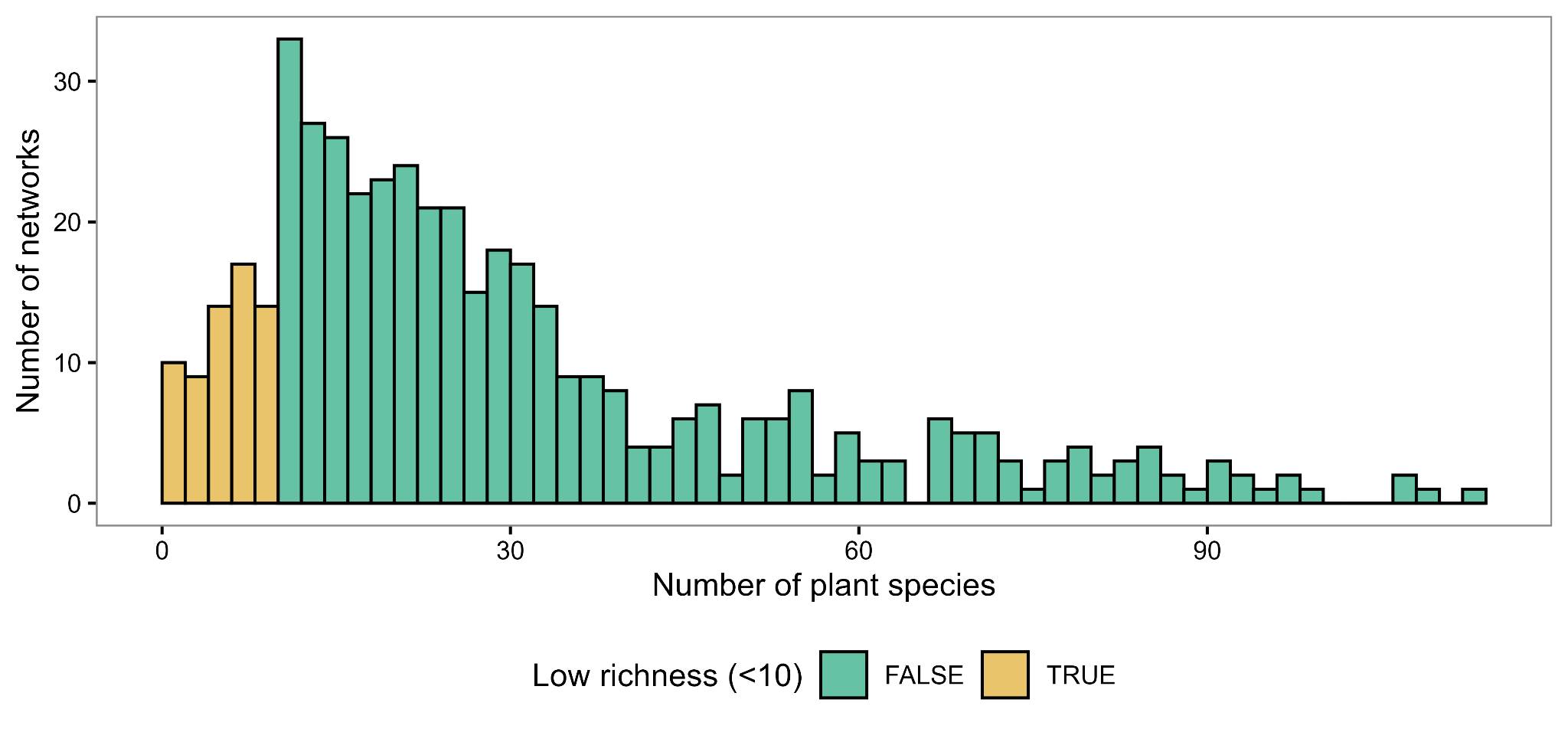

**Supplementary Figure S9.** Distribution of flowering plant species richness in the floral-survey data for networks retained after the interaction-count filter (at least 30 interactions). Plant richness was calculated from non-zero floral-abundance records after excluding missing names and names lacking a species epithet. Networks with fewer than 10 flowering plant species in the floral survey were excluded from the final dataset. Additional richness criteria were applied to the interaction data, requiring at least 10 plant species and 10 pollinator taxa per network.

Networks identified from:

Studies (n = 52)

Networks (n = 1630)

Without separate plant survey

Studies (n = 18)

Networks (n = 595)

With a separate plant survey

Studies (n = 36)

Networks (n = 1035)

Plant species in the survey data largely matched those in the interaction data

Studies (n = 31)

Networks (n = 583)

>25% of plant species, or the most frequently visited species, not represented in the plant survey data

Studies (n = 5)

Networks (n = 452)

Networks with at least 30 interactions

Studies (n = 31)

Networks (n = 459)

**Identification**

**Screening**

Networks with fewer than 30 interactions:

Studies (n = 0)

Networks (n = 124)

Networks with at least 10 plant and pollinator species

Studies (n = 27)

Networks (n = 268)

**Include**

**Included**

Networks with less than 10 plant/ pollinator species

Studies (n = 4)

Networks (n = 191)

**Exclude**

Final studies and networks included in analysis

Studies (n = 27)

Networks (n = 268)

**Supplementary Figure S10.** PRISMA (Preferred Reporting Items for Systematic Reviews and Meta-Analyses) flowchart illustrating the identification, screening, and selection of networks from the EuPPollNet database for inclusion in this study.

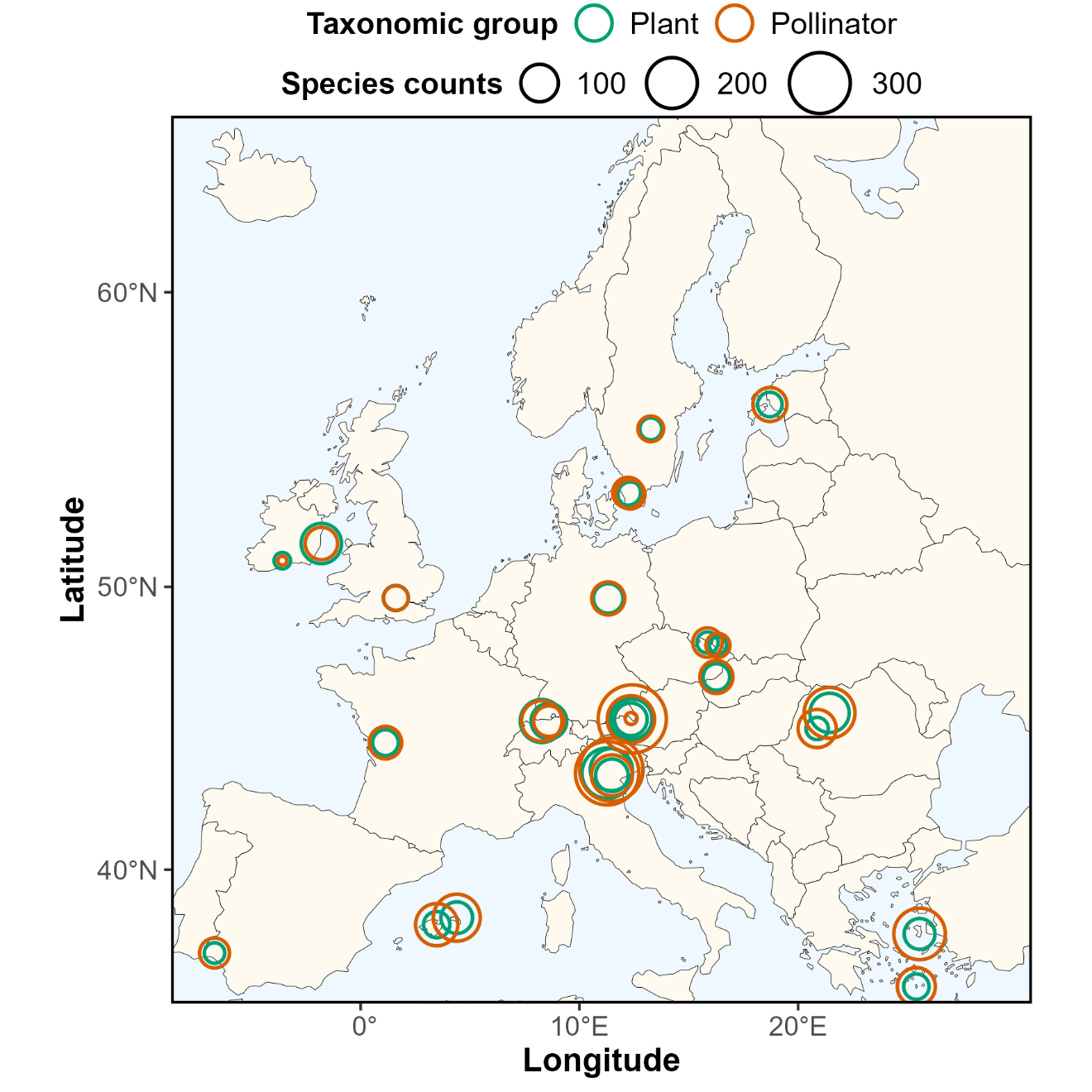

**Supplementary Figure S11.** Map of the 27 study locations across 13 European countries included in the final dataset (268 networks in total). Each location is represented by two open circles indicating the total number of plant species (green) and pollinator species (orange) recorded at the species level across all networks within that study. Circle size is proportional to species richness.

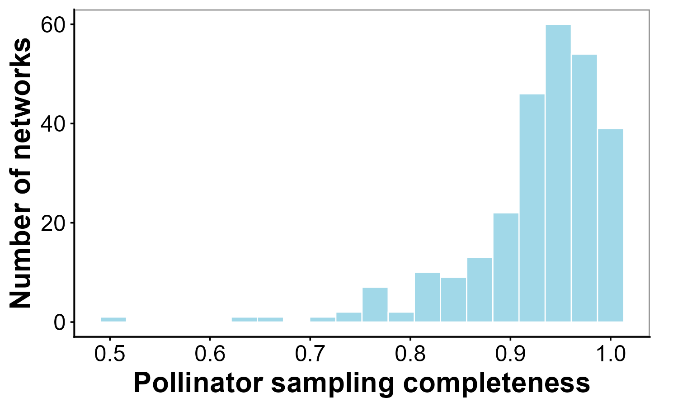

**Supplementary Figure S12.** Histogram of pollinator sampling completeness across the 268 networks retained for considering in this study. Sampling completeness was assessed with abundance-based sample coverage estimator (Ĉ) implemented in the R package iNEXT (Hsieh et al. 2016). Values close to 1 indicate high sampling completeness. Most networks exhibited high coverage, with a median Ĉ of 0.95, indicating that pollinator community diversity was sufficiently sampled in the published networks.

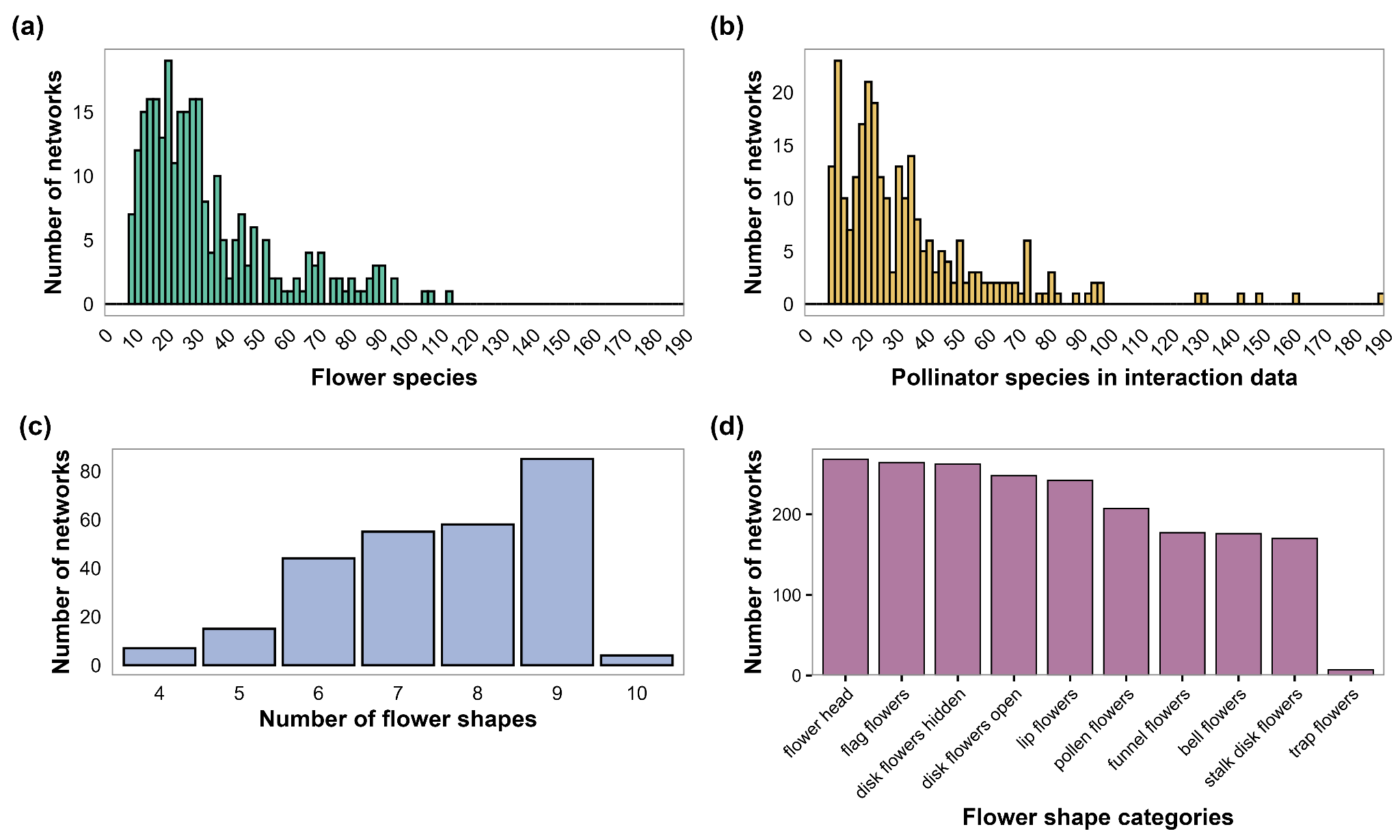

**Supplementary Figure S13.** Distributions of (a) plant species richness, (b) pollinator species richness, and (c) the number of distinct flower-shape categories per network. (d) Number of networks containing each flower-shape category in the floral-survey data.

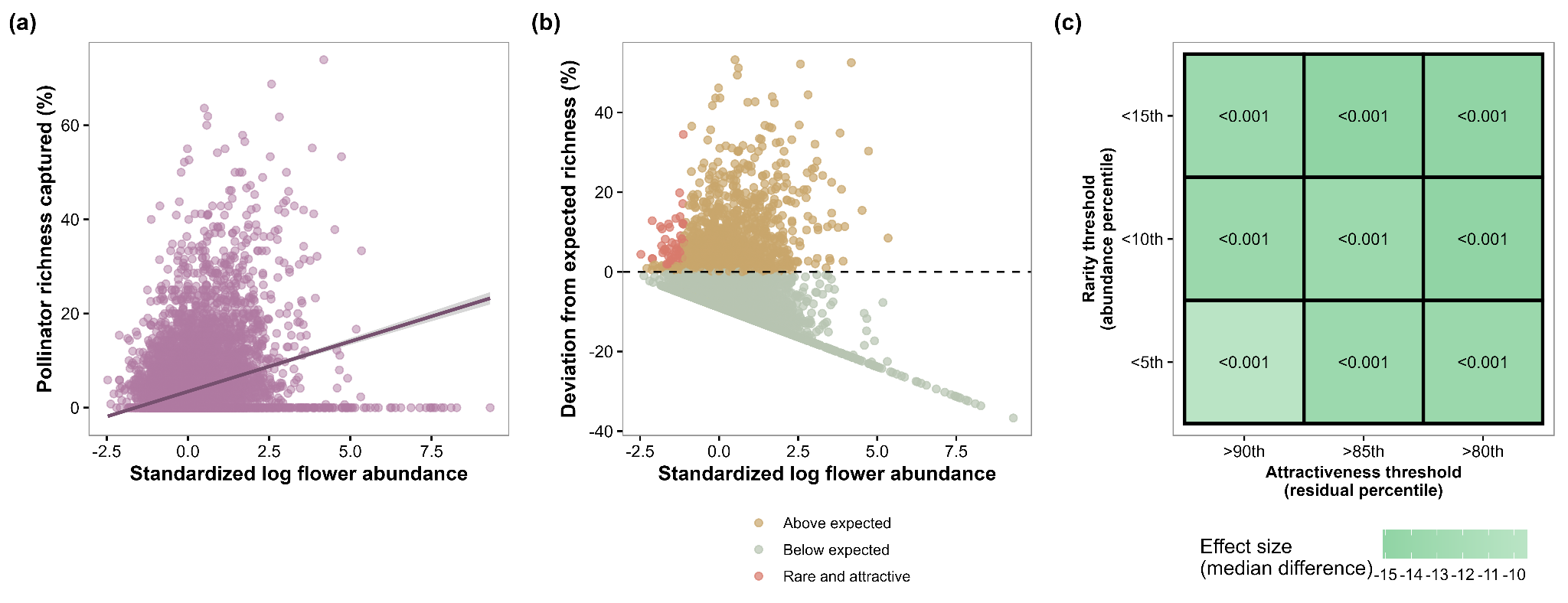

**Supplementary Figure S14.** Identification of rare–attractive plants and sensitivity analysis of their omission from abundance-based subsampling. (a) Relationship between floral abundance and the percentage of network-level pollinator species richness associated with individual plant species. Floral abundance was log-transformed using log(x + 1) and then z-standardized within each network. (b) Deviations between observed richness capture and predictions based on the fixed effects of the linear mixed-effects model. Positive deviations indicate higher-than-expected richness capture given floral abundance. Plant–network observations with standardized log-transformed floral abundance below the 5th percentile and deviations above the 90th percentile were identified as rare–attractive and are highlighted in pink. Both percentile thresholds were calculated across all observations. Species meeting these criteria in at least one network were included in the candidate rare–attractive species list. (c) Sensitivity of the estimated effect of omission to alternative rarity and attractiveness thresholds. For each threshold combination, networks were grouped according to whether selection of the ten most abundant plant species omitted at least one candidate rare–attractive species. The comparison group comprised networks in which no candidate species were present or all candidate species were selected. Effect size is the median percentage of pollinator richness captured in the omission group minus that in the comparison group, expressed in percentage points. Negative values indicate lower richness capture in the omission group. Green indicates negative differences, pink indicates positive differences, and white indicates values close to zero.

**Supplementary Tables**

**Supplementary Table S1.** Pairwise comparisons of the percentage of pollinator richness captured under different plant-subsampling strategies and sampling efforts. Comparisons were conducted using two-sided paired Wilcoxon signed-rank tests across networks with results available under both conditions. Median 1 and Median 2 refer to the median percentage of pollinator richness captured under the first and second conditions listed, respectively. P values were adjusted across all comparisons using the Holm method.

| **Comparison** | **Paired networks (n)** | **Median 1 (%)** | **Median 2 (%)** | **Adjusted P value** |
| --- | --- | --- | --- | --- |
| Top 10 abundant vs Top 5 abundant | 268 | 77.78 | 59.47 | <0.001 |
| Top 5 abundant vs Top 3 abundant | 268 | 59.47 | 42.74 | <0.001 |
| Top 5 flower-shape vs Top 3 flower-shape | 259 | 57.14 | 42.86 | <0.001 |
| Random 10 vs Random 5 | 268 | 49.83 | 29.97 | <0.001 |
| Random 5 vs Random 3 | 268 | 29.97 | 19.36 | <0.001 |
| Top 10 abundant vs Random 10 | 268 | 77.78 | 49.83 | <0.001 |
| Top 5 abundant vs Random 5 | 268 | 59.47 | 29.97 | <0.001 |
| Top 3 abundant vs Random 3 | 268 | 42.74 | 19.36 | <0.001 |
| Top 5 flower-shape vs Random 5 | 261 | 57.14 | 29.84 | <0.001 |
| Top 3 flower-shape vs Random 3 | 265 | 42.86 | 19.46 | <0.001 |
| Top 5 abundant vs Top 5 flower-shape | 261 | 59.38 | 57.14 | 0.036 |
| Top 3 abundant vs Top 3 flower-shape | 265 | 42.86 | 42.86 | 0.749 |
| Top 10 abundant vs Top 10 phylogenetic | 253 | 77.27 | 47.62 | <0.001 |
| Top 5 abundant vs Top 5 phylogenetic | 268 | 59.47 | 26.01 | <0.001 |
| Top 3 abundant vs Top 3 phylogenetic | 268 | 42.74 | 16.23 | <0.001 |

**Supplementary Table S2.** Apid bee species and number of networks in which a species would have been missed if only the ten most abundant plant species were considered in the 268 networks. Species occurring in no more than five networks were excluded to avoid bias from rare occurrences.

| **Pollinator genus** | **Pollinator species** | **Networks total** | **Networks missed** | **Missed rate (%)** |
| --- | --- | --- | --- | --- |
| Amegilla | *Amegilla quadrifasciata* | 6 | 1 | 16.67 |
| Anthophora | *Anthophora furcata* | 7 | 3 | 42.86 |
|  | *Anthophora canescens* | 7 | 2 | 28.57 |
|  | *Anthophora plumipes* | 6 | 1 | 16.67 |
| Apis | *Apis mellifera* | 215 | 11 | 5.12 |
| Bombus | *Bombus campestris* | 12 | 7 | 58.33 |
|  | *Bombus vestalis* | 12 | 7 | 58.33 |
|  | *Bombus rupestris* | 11 | 6 | 54.55 |
|  | *Bombus gerstaeckeri* | 14 | 7 | 50.00 |
|  | *Bombus hypnorum* | 29 | 14 | 48.28 |
|  | *Bombus sylvestris* | 15 | 7 | 46.67 |
|  | *Bombus bohemicus* | 11 | 5 | 45.45 |
|  | *Bombus mendax* | 20 | 8 | 40.00 |
|  | *Bombus humilis* | 50 | 19 | 38.00 |
|  | *Bombus monticola* | 37 | 14 | 37.84 |
|  | *Bombus subterraneus* | 8 | 3 | 37.50 |
|  | *Bombus sylvarum* | 41 | 15 | 36.59 |
|  | *Bombus ruderatus* | 6 | 2 | 33.33 |
|  | *Bombus mucidus* | 16 | 5 | 31.25 |
|  | *Bombus pratorum* | 102 | 31 | 30.39 |
|  | *Bombus jonellus* | 28 | 8 | 28.57 |
|  | *Bombus ruderarius* | 42 | 11 | 26.19 |
|  | *Bombus hortorum* | 112 | 29 | 25.89 |
|  | *Bombus pyrenaeus* | 29 | 7 | 24.14 |
|  | *Bombus argillaceus* | 27 | 5 | 18.52 |
|  | *Bombus terrestris* | 142 | 26 | 18.31 |
|  | *Bombus lucorum* | 55 | 10 | 18.18 |
|  | *Bombus muscorum* | 11 | 2 | 18.18 |
|  | *Bombus wurflenii* | 57 | 9 | 15.79 |
|  | *Bombus lapidarius* | 135 | 21 | 15.56 |
|  | *Bombus mesomelas* | 7 | 1 | 14.29 |
|  | *Bombus pascuorum* | 188 | 26 | 13.83 |
|  | *Bombus barbutellus* | 10 | 1 | 10.00 |
|  | *Bombus soroeensis* | 84 | 6 | 7.14 |
| Ceratina | *Ceratina cyanea* | 20 | 8 | 40.00 |
|  | *Ceratina cucurbitina* | 17 | 6 | 35.29 |
| Eucera | *Eucera nigrescens* | 26 | 11 | 42.31 |
|  | *Eucera oraniensis* | 8 | 2 | 25.00 |
|  | *Eucera longicornis* | 13 | 3 | 23.08 |
| Nomada | *Nomada flavoguttata* | 7 | 6 | 85.71 |
| Xylocopa | *Xylocopa violacea* | 11 | 3 | 27.27 |

**Supplementary Table S3.** Flower abundance sampling methods used across EuPPollNet studies included in this analysis.

| **Study ID** | **Flower sampling methods** |
| --- | --- |
| 2_Petanidou | Number of flowers/sub-inflorescences/stalks/heads/inflorescences/spikes per square meter |
| 4_Marini | Relative abundance of the flowering plant at the site |
| 5_Marini | Relative abundance of the flowering plant at the site |
| 6_Marini | Relative abundance of the flowering plant at the site |
| 7_Scheper | Percentage of flower cover per square centimeter |
| 11_Clough | Number of flowers per 3x3m quadrat |
| 12_Ockinger | Number of flowers per square meter |
| 20_Hoiss | Percentage of flower cover per 4x4m quadrat |
| 22_Kallnik | Percentage of flower cover per square meter |
| 28_Sutter | Percentage of flower cover per square meter |
| 29_Magrach | Percentage of flower cover per square meter |
| 30_Smith | Percentage of flower cover per square meter |
| 31_Roberts | Percentage of flower cover per square meter |
| 36_Larkin | Mean flower number/species per site |
| 37_White | Mean abundance per flowering plant species |
| 38_Maurer | Mean flower number per square meter and site |
| 39_Schweiger | Number of flowers per square meter |
| 40_Knight | Flowers or inflorescences abundance categories (high, medium and low) |
| 41_Knight | Percentage of flowers or inflorescences cover per 60 square meters |
| 42_Knight | Percentage of flowers or inflorescences cover per 60 square meters |
| 44_Knight | Percentage of flower cover |
| 45_Knight | Percentage of flower cover |
| 46_Knight | Percentage of flower cover |
| 47_Benadi | Percentage of flower cover per square meters |
| 50_Hervias-Parejo | Mean flower number per species-site-day |
| 51_Petanidou | Number of flowers per square meter |
| 54_Lazaro | Number of flowers/inflorescences per 30 square meters |

**Supplementary Table S4.** Flower-shape categories based on Kugler (1955, 1970) and the BiolFlor classification. Categories are ordered by the number of distinct plant species recorded in the floral-survey data. Network counts indicate the number of networks with at least one recorded interaction involving plants in each category. Main families are ranked by their frequency among floral-survey records with available family assignments. Example species are selected according to their frequency among positive-interaction records.

| **Flower Shape Category** | **Plant Species (n)** | **Networks (n)** | **Main Families (top 3)** | **Example Species** |
| --- | --- | --- | --- | --- |
| flower head | 210 | 268 | Asteraceae, Caprifoliaceae, Campanulaceae | *Erigeron annuus; Achillea millefolium; Leontodon hispidus* |
| disk flowers hidden | 187 | 240 | Rosaceae, Ranunculaceae, Brassicaceae | *Ranunculus acris; Potentilla reptans* |
| lip flowers | 187 | 218 | Lamiaceae, Plantaginaceae, Orobanchaceae | *Prunella vulgaris; Thymus pulegioides; Betonica alopecuros* |
| disk flowers open | 113 | 213 | Apiaceae, Rubiaceae, Euphorbiaceae | *Daucus carota; Heracleum sphondylium* |
| flag flowers | 102 | 254 | Fabaceae, Polygalaceae | *Trifolium pratense; Lotus corniculatus* |
| bell flowers | 65 | 134 | Campanulaceae, Ericaceae, Primulaceae | *Campanula scheuchzeri;*  *Rhododendron hirsutum* |
| stalk disk flowers | 65 | 121 | Caryophyllaceae, Boraginaceae, Primulaceae | *Petrorhagia saxifraga; Myosotis alpestris* |
| pollen flowers | 63 | 166 | Hypericaceae, Cistaceae, Rosaceae | *Hypericum perforatum;*  *Helianthemum nummularium* |
| funnel flowers | 49 | 136 | Convolvulaceae, Gentianaceae, Caprifoliaceae | *Convolvulus arvensis; Verbena officinalis* |
| trap flowers | 2 | 1 | Apocynaceae | *Vincetoxicum hirundinaria* |

**Supplementary Table S5.** Rare but highly attractive plant species requiring additional sampling.

| Flower shape | Plant species |
| --- | --- |
| bell flowers | *Campanula barbata, Campanula patula* |
| disk flowers hidden | *Capsella bursa-pastoris, Geranium sylvaticum, Oxalis articulata, Ranunculus acris* |
| disk flowers open | *Daucus carota, Peucedanum oreoselinum, Torilis arvensis* |
| flag flowers | *Lotus corniculatus, Trifolium fragiferum, Trifolium pratense, Vicia sepium* |
| flower head | *Achillea millefolium, Astrantia major, Carduus acanthoides, Carduus defloratus, Centaurea jacea, Centaurea nigra, Centaurea scabiosa, Centaurea stoebe, Cirsium arvense, Cirsium vulgare, Crepis aurea, Crepis capillaris, Crepis setosa, Erigeron annuus, Eryngium maritimum, Helianthus annuus, Helminthotheca echioides, Hieracium umbellatum, Hypochaeris radicata, Knautia arvensis, Lapsana communis, Leontodon hispidus, Matricaria chamomilla, Picris hieracioides, Taraxacum campylodes, Taraxacum officinale* |
| lip flowers | *Echium vulgare, Iris pseudacorus, Lamium album, Odontites vernus, Rhinanthus minor* |
| pollen flowers | *Hypericum androsaemum, Hypericum perforatum, Papaver rhoeas* |
